# Phosphite, an analog of phosphate, counteracts Phosphate Induced Susceptibility of rice to the blast fungus *Magnaporthe oryzae*

**DOI:** 10.64898/2026.01.22.700763

**Authors:** Mani Deepika Mallavarapu, Héctor Martín-Cardoso, Gerrit Bücker, Melissa Alussi, Antoni García-Molina, Blanca San Segundo

**Author notes:** Correspondence: Blanca San Segundo,; Antoni García-Molina. E-mail address Mani Deepika Mallavarapu,. Héctor Martín-Cardoso, Gerrit Bücker, Melissa Alussi, Antoni Garcia-Molina, Blanca San Segundo.

## Abstract

Phosphate (Pi) and phosphite (Phi), a non-metabolizable analogue of Pi, are taken up by plant roots through the same transport system. Whereas Pi is an essential nutrient for plants, Phi might function as a biostimulant and in protection against pathogens. However, how Phi mechanistically exerts beneficial effects on plants remains unsolved. We examined the impact of Phi and Pi on *Arabidopsis thaliana* and rice growth and upon pathogen infection. Phi inhibited the *in vitro* growth of *Plectosphaerella cucumerina* and *Fusarium fujikuroi* in a dose-dependent manner, whereas *Magnaporthe oryzae* growth was largely unaffected. Phi’s effect on plant growth was dependent on the plant species, the basal Pi level in the plant, and the ratio Pi to Phi. In Arabidopsis, Phi enhanced resistance to *P. cucumerina* by triggering a hypersensitive response-like cell death. Notably, Phi reversed Pi-induced susceptibility to blast (*M. oryzae*) and bakanae (*F. fujikuroi*) diseases in rice. Transcriptomic analysis revealed that Phi triggered extensive reprogramming in rice under high Pi, including the activation of signaling pathways enriched in phosphorylation-dependent processes, while attenuating induction of carbon metabolism. Phi acts as a multifaceted agent, promotes balanced metabolic state, improved plant performance, and reduced Pi-induced disease susceptibility when applied under appropriate Pi conditions.

**Highlight:** Phosphite application confers protection against fungal pathogens in Arabidopsis and rice plants by regulating signaling pathways depending on phosphorylation processes.

## Introduction

Phosphorus (P) is an essential element required for plant growth and development. It is a component of key molecules, like nucleic acids, phospholipids and adenosine triphosphate (ATP), the main energy currency of the cell. P is also a central regulator in numerous metabolic reactions and signaling pathways, and modulation of protein activity through phosphorylation (Wang *et al*., 2021). Plants take up P primarily through the root system, mainly as inorganic phosphate (Pi), which is then metabolized by the plant. Although P levels in soils are generally high, its bioavailability is often extremely low due to absorption and precipitation reactions involving soil minerals (e.g., iron, aluminum, calcium) (Hinsinger *et al*., 2011). To overcome Pi limitation, P fertilizers are routinely used to maintain optimal productivity in agricultural soils. Yet, the heavy dependence on chemical fertilizers has become increasingly problematic, driving economic, ecological, and sustainability concerns. Overapplication of Pi fertilizers frequently results in environmental contamination, eutrophication of aquatic ecosystems, and accelerated depletion of finite global phosphate reserves (Herrera-Estrella and López-Arredondo, 2016). Given these critical environmental and Pi resource limitations, there is a pressing need to explore alternative strategies to optimize Pi nutrition while maintaining crop yields and mitigating adverse environmental impacts of chemical fertilizers.

Beyond being an essential plant nutrient, evidence supports that Pi plays a fundamental role in regulating plant immunity. The relationship between Pi and disease resistance in plants is, however, complex as Pi’s effect appears to depend on the plant species and the type of pathogen (Chan *et al*., 2021). Studies so far carried out indicate that high Pi fertilization can promote either resistance or susceptibility to pathogen infection, as illustrated by studies conducted in *Arabidopsis thaliana* and rice (*Oryza sativa*) plants. In rice, elevated Pi levels enhance susceptibility to infection by the fungal pathogens *Magnaporthe oryzae*, and *Fusarium fujikuroi*, the causal agents of blast and bakanae disease, respectively (Campos-Soriano *et al*., 2020; Martín-Cardoso *et al*., 2025). Contrary to what is observed in rice, Arabidopsis plants accumulating Pi showed increased resistance to fungal pathogens, such as *Plectosphaerella cucumerina* and *Colletotrichum higginsianum* (Val-Torregrosa *et al*., 2022*b*). Additionally, Pi supply has been shown to modulate the expression of immune responses in rice and Arabidopsis (Castrillo *et al*., 2017; Campos-Soriano *et al*., 2020; Val-Torregrosa *et al*., 2022*a*,*b*; Martín-Cardoso *et al*., 2024). Further illustrating the complexity of the impact of Pi nutrition in disease resistance, very recently, it has been reported that certain fungal pathogens secrete effectors into the host plant that hijack host Pi phosphate sensing pathways to suppress immunity, thereby facilitating disease susceptibility (McCombe *et al*., 2025). At the molecular level, protein phosphorylation events serve as central regulators of plant immune responses against pathogen infection (Hake and Romeis, 2019; Erickson *et al*., 2022). Central to this process is that pathogen recognition triggers the induction of protein kinase signaling cascades for the activation of multiple regulatory networks leading to defense manifestation. The integration of phosphorylation signaling cascades might exert a positive or negative control of the plant immune system, both at the local and systemic level. Altogether, these findings underscore the intricate interplay between Pi nutrition and disease resistance. However, the exact mechanisms by which Pi signaling interacts with immune signaling remain largely unknown.

Unlike Pi, phosphite (Phi), a reduced form of phosphate, is highly soluble, and readily absorbed by plants via phosphate transporters. However, plants cannot metabolize Phi due to lack of enzymes required for converting Phi into the usable form Pi. The application of Phi has been reported as a strategy to effectively control weed species, which are resistant to herbicide treatments (Achary *et al*., 2017). Importantly, during the past years, Phi’s utility as a compound effective for the control of diseases such as those caused by oomycetes and fungal pathogens has gained much attention (Achary *et al*., 2017; Havlin and Schlegel, 2021; Mohammadi *et al*., 2021; Li *et al*., 2025). A dual mode of action has been proposed to explain the inhibitory activity of Phi on the growth of phytopathogens by either direct inhibition of fungal growth, or by triggering the plant’s own defense mechanisms (Massoud *et al*., 2012; Mehta *et al*., 2022). Furthermore, Phi’s application appears to protect certain plant species either by soil application (e.g., fertigation), or by foliar application (Havlin and Schlegel, 2021; Mohammadi *et al*., 2021). In particular, foliar spray of Phi has proven to be effective in reducing blast disease severity in the *indica* rice cv. BPT5204 (a widely cultivated rice variety in India) (Mehta *et al*., 2022).

Phi’s effects on plant growth are expected to be intertwined with plant nutrition, particularly with Pi nutrition. In Arabidopsis, Phi has been shown to selectively inhibit Phosphate Starvation Responses (PSRs), a survival strategy that plants deploy when they lack sufficient Pi in the soil (Ticconi *et al*., 2001; Jost *et al*., 2015; Pérez-Zavala *et al*., 2024). At present, however, the underlying molecular mechanisms of Phi’s protective effect and precise interactions between Phi and plant immunity in rice remain to be fully elucidated.

Given the complex behavior of Phi in plant nutrition and disease resistance, understanding its role is critical for agricultural applications from an ecological perspective for sustainable crop production. For this, several critical gaps persist in our understanding on beneficial effects that plants can receive from Phi application, especially in crop species. Firstly, although Phi can influence both growth and immunity, the trade-offs between Phi-induced growth modulation and defense activation have not been systematically explored. Secondly, while Phi treatment can protect plants from diseases, the molecular mechanisms underlying Phi-mediated modulation of plant immune responses remain poorly defined. Thirdly, plant responses to Phi appear to vary significantly across species, yet comparative studies examining monocots like rice and dicots like Arabidopsis are limited. These studies are essential to identify divergent and conserved evolutionary solutions underpinning adaptive mechanisms to Pi supply and/or Phi application among species. Lastly, the impact of Phi application in the context of current agricultural practices based on repeated use of Pi fertilizers remains largely uninvestigated. A better knowledge of trade-offs between growth and defense activation upon Phi application under different Pi fertilization regimes is particularly important as it may affect overall plant fitness and disease resistance.

Based on these critical knowledge gaps, this study aimed to evaluate the effects of Phi application in growth and disease resistance in Arabidopsis and rice plants, the two plant species adopted as models for studies in monocotyledonous and dicotyledonous species, respectively. The inhibitory activity of Phi on the growth of phytopathogens that are responsible for significant economic losses in dicotyledonous plants (e.g., *Plectosphaerella cucumerina*) and monocotyledonous plants (e.g., *Magnaporthe oryzae* and *Fusarium fujikuroi*) has been investigated. Furthermore, knowing that Pi influences disease resistance in Arabidopsis and rice plants in a differential manner (e.g., high Pi supply confers resistance and susceptibility to pathogen infection in Arabidopsis and rice plants, respectively), the effect of Phi in disease resistance in Arabidopsis and rice plants has been examined. Importantly, the effect of Phi in Arabidopsis and rice growth and disease resistance was evaluated in plants that have been grown under different Pi and/or Phi regimes. Special emphasis was placed on rice, a staple food for over half the world’s population, and a crop of importance in global food security. Here, we investigated whether the application of Phi can counteract Pi-induced susceptibility to infection by *M. oryzae* in rice plants. Finally, we systematically investigated transcriptional reprogramming triggered by Phi application in rice. We show that Phi application caused reprogramming of genes involved in phosphorylation signaling, ethylene signaling, and nutrient transport. The information gained in these studies provides a more holistic understanding of Phi’s function in plants. In rice, a Phi-induced priming contributing to attenuate Pi-induced blast susceptibility is proposed. Together, this study provides a foundation for developing more sustainable strategies for rice production that rely less on high Pi fertilization and chemical fungicides.

## Materials and Methods

### Plant materials and growth conditions

*Arabidopsis thaliana* (ecotype Columbia-0) seeds were surface-sterilized in 1% sodium hypochlorite containing 0.05% SDS by gently shaking for 8–10 minutes, extensively rinsed with sterile water (3 times), and air-dried. Seeds were stratified at 4°C in the dark for 3–4 days. Seedlings were initially grown for one week on ½ MS medium to establish uniform growth and then transferred to modified half-strength Hoagland’s medium containing varying concentrations of potassium phosphate (Pi, KH_2_PO_4_, Merck KGaA, Darmstadt, Germany) and/or potassium phosphite (Phi, KH_2_PO_3_, Biosynth, Cymit Química S.L. Spain) for one additional week. Arabidopsis plants were grown under controlled conditions with an 8-hour light/16-hour dark photoperiod, at 22°C, with light intensity of 110 µmol mC² sC¹ and 60% relative humidity.

*Oryza sativa* (Nipponbare cv, *japonica* subspecies) seeds were heat-treated (at 37°C) in the dark for 3–4 days to promote uniform germination. The rice cultivars Argila, Baixet, Bomba and Guara (*O. sativa, japonica* varieties) were also used in infection experiments. Rice plants were grown on substrate mix composed of 50% inert base (33% vermiculite and 66% Turba Floratorf® peat supplemented with 1 g/L CaCOC) and 50% DORSILIT^®^ quartz sand (0.6–1.2 mm), under controlled greenhouse conditions with a 14-hour light/10-hour dark photoperiod, at 28°C and 60% relative humidity. During the first week, plants were watered, followed by weekly fertigation with half-strength modified Hoagland’s solution with the appropriate Pi and Phi concentrations for the next two weeks.

Three different Pi concentrations were used to grow Arabidopsis and rice plants. They were: 0.025mM Pi, 0.25mM Pi, and 2.5mM Pi (representing low, optimal, and high Pi conditions, respectively) (Campos-Soriano *et al*., 2020). To assess the effect of Phi, different Pi and Pi+Phi ratios were tested initially (Supplementary Fig. S1). After experimental trials, Phi at 1/10th ratios of the respective Pi concentration (e.g., 0.1x concentrations in Supplementary Fig. S1) were selected as the standard condition for further experiments, as follows: 0.025mM Pi + 0.0025mM Phi, 0.25mM Pi + 0.025mM Phi, and 2.5mM Pi + 0.25mM Phi.

### Fungal material

*Plectosphaerella cucumerina* and *Fusarium fujikuroi* (297 isolate) were grown on PDA (Potato Dextrose Agar), while *Magnaporthe oryzae* (strain Guy11) was grown on CMA (Complete Media Agar), containing 30mg/L chloramphenicol. Fungi were grown in their respective media for about 2 weeks at 25°C under a 16h light/8h dark cycle. Fungal spores were collected from each plate by adding sterile water to the mycelium surface, scraping and filtering the spore suspension using Miracloth (Merck KGaA, Darmstadt, Germany). The spore solution was adjusted to the desired final concentration using a Neubauer Chamber (5×10^5^ spores/mL for *M. oryzae* and *P. cucumerina*; 1×10^6^ spores/mL for *F. fujikuroi*). For infection experiments with *M. oryzae*, 0.02% Tween^®^ 20 (MilliporeSigma) was added to the spore suspension.

### Effect of Pi and Phi on *in vitro* growth of fungal pathogens

The effect of Pi or Phi on the *in vitro* growth of fungal pathogens (e.g., *P. cucumerina*, *M. oryzae* and *F. fujikuroi*) was determined using a microtiter plate assay previously described (Cavallarin *et al*., 2007). For this, fungi were grown in 96-well microtiter plates with respective media (150µL, containing chloramphenicol 30mg/L) at 25°C in the dark. Fungal spores (50µL of 5×10^5^ spores/mL) were added to microtiter wells. The effect of Phi or Pi on fungal growth was determined by measuring the absorbance at 595nm of fungal cultures with time (up to 48h) using SpectraMax^®^ M3 plate reader (Molecular Devices LLC, Sunnyvale, CA, USA).

### Infection assays

For infection experiments in *Arabidopsis thaliana*, 2-week-old *in vitro*-grown seedlings were spray-inoculated with a *P. cucumerina* spore suspension (5 × 10C spores/mL), or mock-inoculated using an atomizer (two pumps per plant) as previously described (Val-Torregrosa *et al*., 2022*b*). Following inoculation, seedlings were maintained under the same growth conditions and disease symptoms were assessed at 7 days post-inoculation (dpi). Three independent experiments were conducted, each with a minimum of 24 plants per treatment.

For rice infection assays, plants at the 3-4 leaf developmental stage were spray-inoculated with a *M. oryzae* spore suspension (5 × 10C spores/mL), or mock-inoculated, using an aerograph at 2 atm pressure (2–3mL per pot, with 10-12 plants per pot). Rice plants were kept overnight in darkness and high humidity using plastic bags and allowed to continue growth. Infection progression was monitored over time, and visible symptoms were documented at 7 days post-inoculation (dpi). The percentage of leaf area affected by blast lesions was quantified using APS Assess 2.0 software (Lamari, 2008). Three independent experiments were performed with at least 24 plants per condition in each experiment.

Infections with *F. fujikuroi* were carried out as described (Martín-Cardoso *et al*., 2025). Briefly, rice seeds were pregerminated on petri dishes under Pi or Pi+Phi supply and then inoculated with *F. fujikuroi* spores (1×10^6^ spores/mL) and kept in the dark for 24h. After the infection and transplant to soil, seedlings were supplied with half-strength Hoagland’s solution at the appropriate Pi or Pi+Phi concentration twice per week. Measurements of root length were carried out by image analysis at 10 days post-inoculation (dpi) using ImageJ/Fiji v1.54j software (http://fiji.sc/Fiji). The severity of disease symptoms was estimated at 10 days after infection for each Pi and Pi+Phi condition as described (Amatulli *et al*., 2010), using the following scale: 0, symptomless seedlings; 20, stunted seedlings with yellow and narrow leaves; 40, thin seedlings with pale-yellow and narrow leaves; 60, seedlings with pale-yellow and narrow leaves, long and thin internodes and short roots; 80, seedlings with some necrosis of the crown and severe bakanae symptoms; 100, dead seedlings. Three independent experiments were carried out (N=15 per condition, each experiment).

For quantification of fungal biomass, genomic DNA was extracted from infected leaves using MATAB extraction method (0.1 M Tris-HCl pH 8.0, 1.4 M NaCl, 20 mM EDTA, 2% MATAB, 1% PEG 6000 and 0.5% Na_2_SO_3_) (Murray and Thompson, 1980). Fungal biomass was quantified by quantitative PCR (qPCR) using specific primers for the corresponding fungus. qPCR primers are listed in Supplementary Table S1, designed for detection of *P. cucumerina* β-tubulin, *M. oryzae* 28S rDNA and *F. fujikuroi* 28S rDNA (Campos-Soriano *et al*., 2020; Val-Torregrosa *et al*., 2022*b*; Martín-Cardoso *et al*., 2025). The *Ubiquitin21* and *Ubiquitin1* genes were used as the internal control for quantification of fungal biomass in Arabidopsis and rice, respectively.

### Staining methods

For trypan-blue staining of *P. cucumerina*-infected Arabidopsis plants, leaves were fixed by vacuum infiltration for 1 hour in ethanol:formaldehyde:acetic acid (80:3.5:5 v/v), stained with trypan blue solution for 1 hour, and washed with 70% ethanol three times. The stained leaves were mounted on glass slides with glycerol and observed under bright-field illumination on a Leica DM6 microscope (Leica Microsystems, Wetzlar, Germany).

For staining fungal hyphae growing in liquid medium, Calcofluor White (Sigma) solution (10µL; 1µg/mL) was added to 10µL of fungal cultures from microtiter plate assays and observed under UV light illumination on a Leica DM6 microscope. For UV imaging, excitation and emission wavelengths were preset to 340–380 nm and 425 nm, respectively. Images were analyzed using ImageJ software.

### Measurement of Photosynthetic Pigment content

Chlorophyll content of Arabidopsis was estimated spectrophotometrically based on absorbance of pigment extracts, following the method of Lichtenthaler and Buschmann, 2001. Briefly, leaf samples (10 mg) were flash-frozen in liquid nitrogen and homogenized on a TissueLyser II (QIAGEN, Hilden, Germany). Photosynthetic pigments were extracted in 96% (v/v) ethanol, and absorbance was measured using a SpectraMax® M3 plate reader at 649 nm and 665 nm for chlorophylls, and at 470 nm for carotenoids. Pigment concentrations were calculated using standard equations and expressed in mg/g fresh weight (FW). Three independent experiments were conducted, each with four biological replicates per condition, consisting of six pooled plants per replicate.

Chlorophyll content of rice leaves was estimated non-destructively using a SPAD-502 (SPAD ‘Soil Plant Analysis Development’, Konica Minolta, Japan) chlorophyll meter. For measurements, fully developed leaves (3^rd^ or 4^th^ leaf from 3-4 leaf developmental stage plants) were used, ensuring uniform sampling across treatments. SPAD readings were taken at three positions per leaf: the base, middle, and tip to account for variability. An average of 10-15 readings per plant was recorded to ensure accuracy and reproducibility.

### RNA extraction and gene expression analysis

Total RNA was isolated using Maxwell RSC Plant RNA Kit (Promega). For Reverse transcription (RT)-qPCR, total RNA (1μg) was used for first-strand cDNA synthesis using High Capacity cDNA Reverse Transcription Kit (Applied Biosystems, Waltham, MA, USA). Primers were designed using the NCBI’s Primer-BLAST tool. RT-qPCR reactions were done using SYBR^®^ green (Roche) in optical 384-well plates and the Light Cycler 480 (Roche). In Arabidopsis, Ct values were normalized to *Ubiquitin21* gene (At5g25760). In rice, Ct values were normalized to *Ubiquitin1* gene (Os06g0681400). Four biological replicates and two technical replicates were analyzed for each biological replicate, consisting of a pool of 6 plants. Primers are listed in Supplementary Table S1.

### Phosphate quantification

Free inorganic phosphate (Pi) content in plant tissues was estimated using a colorimetric assay based on molybdenum blue complex formation (Ames, 1966; Versaw and Harrison, 2002). Tissue samples were flash-frozen in liquid nitrogen and homogenized on a TissueLyser II. Pi was extracted using 1% glacial acetic acid (1mL/10mg of tissue, in 2mL Eppendorf tubes). Then, a reaction mix containing 0.7mL of a 1:6 solution of 10% ascorbic acid and 0.42% ammonium molybdate in 1N HCSOC was added to 0.3mL of the sample (1:3 dilution). The resulting blue-colored complex, formed upon reaction with free POC³C, was measured at 820nm using a SpectraMax^®^ M3 plate reader. Free Pi content was quantified based on absorbance values using a standard phosphate calibration curve.

### RNA sequencing and Data analysis

Total RNA was obtained from leaves of rice plants grown under high Pi condition (e.g., 2.5mM Pi), with or without Phi supplementation (N=3 biological replicates). Concentration and purity of RNA samples were assessed using a Nanodrop2000 spectrophotometer (Thermo Scientific), while RNA integrity was confirmed using Agilent 2100 and LabChip GX (Perkin Elmer). RNA sequencing (RNA-Seq) was carried out by Sequentia Biotech (Barcelona, Spain) using the Illumina NovaSeq 6000 platform with PE150 chemistry. RNA-Seq datasets were analysed using a Salmon-DESeq2 pipeline (Love *et al*., 2014; Patro *et al*., 2017) based on the *Oryza sativa* subsp. *japonica* reference genome (IRGSP-1.0). Differentially expressed genes (DEG) were declared for transcripts displaying significant changes in pair-wise comparisons (adjusted *P* ≤ 0.05, Wald’s test). Significant enrichment in Gene Ontology terms for biological processes was computed in *Panther* (https://pantherdb.org) (fold-enrichment ≥ 2, adjusted *P* ≤ 0.05, Fisher’s exact test).

### Statistical analyses

The statistical methods used are described in the respective figure legends (Student’s *t*-test, one-way ANOVA and two-way ANOVA followed by Tukey’s multiple comparison test). Statistical calculations were performed using at least three independent biological replicates (see figure legends for details). Data was analyzed using GraphPad Prism v10.4.1 software (https://www.graphpad.com/).

## Results

### Effect of phosphite and phosphate on growth of the phytopathogenic fungi *P. cucumerina*, *M. oryzae* and *F. fujikuroi*

We investigated whether Phi has inhibitory activity on the growth of important fungal plant pathogens, namely *P. cucumerina*, *M. oryzae* and *F. fujikuroi*. The fungus *P. cucumerina* (formerly known as *Fusarium tabacinum*) is a soil-born pathogen that has worldwide distribution and wide plant-host range. It causes blight and sudden death disease in many dicotyledonous species, including members of the *Brassicaceae* family, and the model plant *A. thaliana* (Ramos *et al*., 2013). *M. oryzae* affects several cereal species, especially rice. It causes rice blast, one of the most destructive diseases for cultivated rice worldwide (Wilson and Talbot, 2009). As for *F. fukikuroi*, this is the causal agent of bakanae (‘foolish seedling’) disease whose incidence has progressively increased in major rice-producing regions (Prà *et al*., 2010).

To assess the effect of Phi on fungal growth, the fungus was grown in liquid cultures in the absence of Phi (control cultures) or in the presence of increasing concentrations of Phi (0.1mM Phi, 0.5mM Phi, 0.75mM Phi and 1mM Phi). The absorbance of fungal cultures was measured over time. As shown in **Fig. 1A** (upper panel), Phi was found to restrict *P. cucumerina* growth by 48h of incubation, at a concentration of 0.5 mM and above, its effect further increasing at higher Phi concentrations. Similarly, growth assays on solid media supplemented with increasing concentrations of Phi revealed inhibition of *P. cucumerina* mycelial growth by Phi (Supplementary Fig. S2). In contrast, Phi did not inhibit the *M. oryzae* growth in liquid culture (**Fig. 1A**, middle panel). Similar to what was observed in *P. cucumerina* cultures, *F. fujikuroi* growth was significantly reduced in the presence of Phi (**Fig. 1A**, lower panel). Calcofluor White staining of fungal cultures confirmed the inhibitory activity of Phi on *P. cucumerina* and *F. fujikuroi* growth (**Fig. 1B**). The presence of Phi at increasing concentrations progressively reduced hyphal growth and branching in *P. cucumerina* and *F. fujikuroi* cultures, while long and highly branched hyphae were observed in control cultures (0mM Phi) (**Fig. 1B**).

**Fig. 1.**
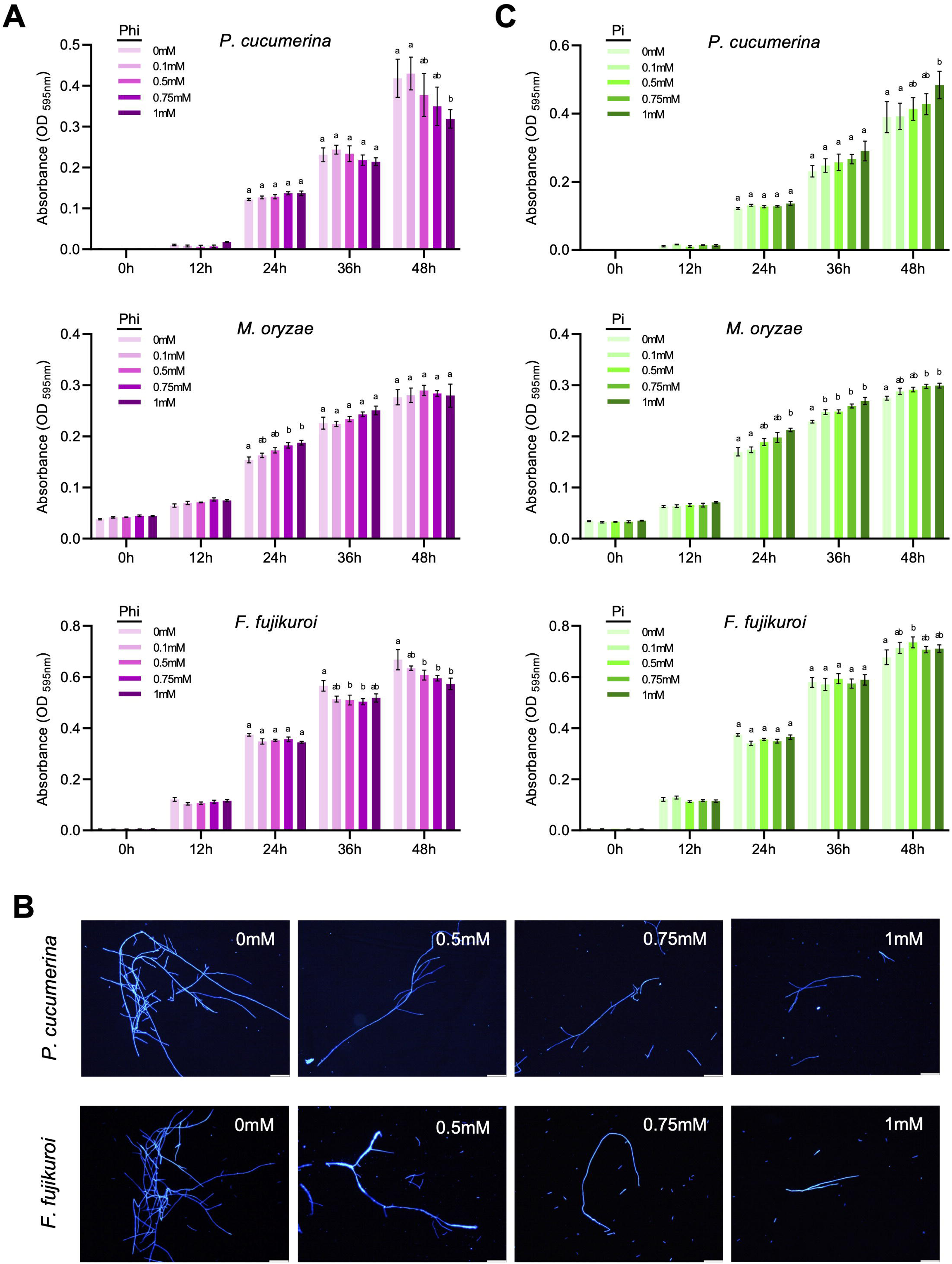
Effect of Phi and Pi on the *in vitro* growth of the plant fungal pathogens *P. cucumerina*, *M. oryzae* and *F. fujikuroi*. (**A and C**) Growth of fungal cultures was determined by measurement of absorbance at 595nm at different times after the addition of either Phi (**A**) or Pi (**C**) at the indicated concentrations. Control cultures were not supplemented with Phi (0mM Phi). Data represents mean ± SEM of 3 biological replicates. Letters above the bars indicate statistically significant differences among treatments according to one-way ANOVA followed by Tukey’s multiple comparison test (*p* < 0.05). Bars sharing the same letter are not significantly different compared to control. (**B**) Fluorescence microscopy images of calcofluor white-stained fungal cultures at 48h of growth with increasing Phi concentrations. Scale bars = 100µm.

Parallelly, we also examined the influence of Pi supplementation on fungal growth. In the presence of increasing concentrations of Pi, *P. cucumerina* growth increased (**Fig. 1C**, upper panel). At a lesser extent, *M. oryzae* growth also increased by incubation with Pi (**Fig. 1C**, middle panel). In contrast, *F. fujikuroi* growth was not as substantially affected by increasing the addition of Pi to the medium (**Fig. 1C**, lower panel).

Together these observations indicated that Phi and Pi have an effect on the *in vitro* growth of fungal pathogens, and that this effect depends on the specific fungal pathogen under consideration. Phi inhibits *P. cucumerina* and *F. fujikuroi* growth in a dose-dependent manner, while it had an insignificant effect on *M. oryzae* growth at the assayed concentrations. Compared with Phi, Pi supplementation had different effects on fungal growth. Pi exerted a stimulatory effect on *P. cucumerina* and *M. oryzae* growth, but it did not affect *F. fujikuroi* growth.

### Effect of phosphite on the growth of Arabidopsis and rice plants

To assess the effect of Phi on the growth of *Arabidopsis thaliana* (ecotype Columbia-0), two sets of plants were grown in parallel. One set of plants was supplied with only Pi (Pi condition), while the other set was supplied with a combination of Pi and Phi (+Phi condition). Three Pi regimes were assayed, namely 0.025mM Pi (henceforth P_0.025_), 0.25 mM Pi (P_0.25_), and 2.5mM Pi (P_2.5_). For each Pi condition, the Phi concentration used was one tenth the Pi concentrations (e.g., 0.025mM Pi + 0.0025mM Phi, 0.25mM Pi + 0.025mM Phi, and Pi 2.5mM Pi + 0.25mM Phi). Plants were grown for one week on half-strength MS medium and for two weeks more under the corresponding Pi or Pi + Phi regime.

In the absence of Phi supply, no differences in plant growth across Pi conditions were observed (**Fig. 2A, B**, upper panels), which was confirmed by fresh weight measurements (**Fig. 2C**, left panel-green bars). However, the content of photosynthetic pigments, both chlorophylls and carotenoids, decreased with increasing Pi supply (without Phi supplementation) (**Fig. 2C**, middle and right panels-green bars). Notably, striking changes were seen in plant growth and photosynthetic pigment content upon Phi addition depending on the Pi condition. At the lowest Pi concentration (P_0.025_ plants) Phi supplementation severely affected plant growth compared with controls with only Pi (**Fig. 2A, B**, lower panels), which was accompanied by a marked reduction in fresh weight, chlorophyll and carotenoid content (**Fig. 2C**, P_0.025_ - pink bars). These observations suggest that Phi either exacerbates Pi deficiency symptoms or has a toxic effect on Arabidopsis plants under limiting Pi conditions. However, under optimal and high Pi conditions (P_0.25_ and P_2.5_ plants, respectively), Phi supplementation significantly increased fresh weight, its effect being greater in P_0.25_ plants than in P_2.5_ plants (**Fig. 2A-C**). Furthermore, P_0.25_ plants supplemented with Phi exhibited greener leaves and accumulated more photosynthetic pigments than plants grown with only Pi (**Fig. 2B, C**). Collectively these results indicated that Phi could act as a biostimulant of Arabidopsis growth when the plants are grown under optimal Pi conditions.

**Fig. 2.**
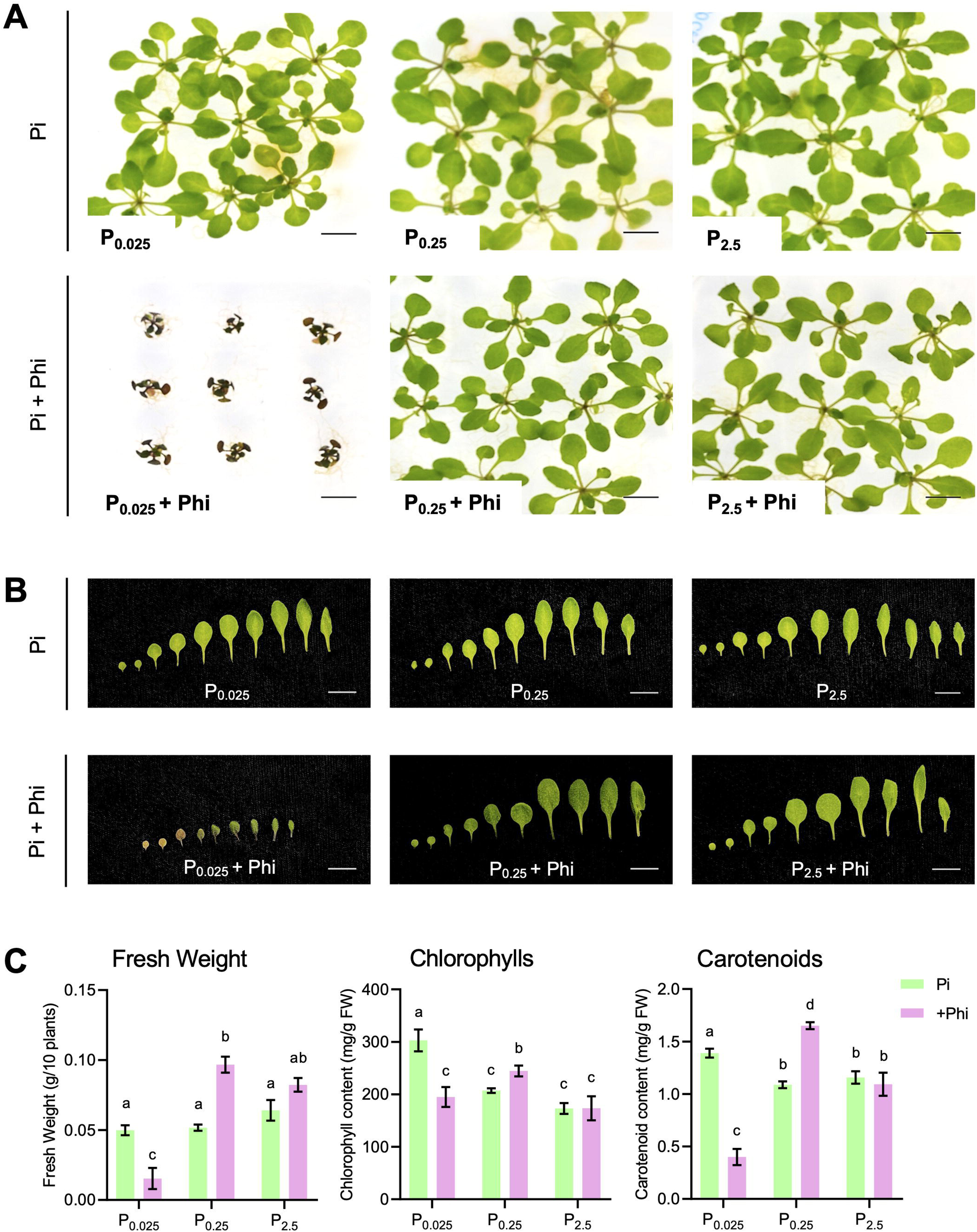
Phosphite supplementation influences growth and photosynthetic pigment content in *A. thaliana*. Plants were grown for 1 week on half-strength MS and then under different Pi regimes (0.025mM Pi, 0.25mM Pi, or 2.5mM Pi) for two weeks, with or without Phi supplementation in each Pi condition (at a 1/10^th^ Phi/Pi molar ratio in all Pi conditions). (**A**) Representative images of *Arabidopsis* plants under each Pi and Pi + Phi condition. (**B**) Leaf series from plants shown in B reflecting differences in the size and pigmentation. Scale bars in B and C, correspond to 10 mm. (**C**) Fresh weight, total chlorophyll content and carotenoid content of plants under each condition. Bars represent ± SEM of at least 4 biological replicates (6 plants per replicate). Letters above bars indicate statistical significance according to two-way ANOVA and Tukey’s post-hoc tests, within and between the Pi and Pi+Phi treatments. Same letters indicate no statistical difference.

Next, we investigated the effect of Phi on rice growth. For this, rice (*O. sativa* cv. Nipponbare) seedlings were grown with water for 1 week and then supplied with P_0.025_, P_0.25_, or P_2.5_, either with or without co-treatment with Phi as one-tenth of P. Phi supplementation did not alter plant fitness under P_0.025_ or P_2.5_, while in plants under P_0.25_, plants shoot length and fresh weight slightly increased (**Fig. 3A, B**). SPAD measurements revealed that Phi reverted the loss in chlorophyll at P_2.5_ due to excess-Pi stress to levels almost comparable to those in P_0.25_ plants. Notably, co-treatments with higher Phi doses – up to 0.25 and 2.5 mM in P_0.25_ and P_2.5_, respectively - were not harmful for plants, but 25 mM Phi negatively impacted in fresh weight and returned to low chlorophyll content in P_2.5_ plants (**Fig. 3**), suggesting Phi phytotoxicity at very high concentrations. Together, these studies demonstrated a differential effect of Phi application on Arabidopsis and rice growth, its application also showing dosage-dependent and Pi-context specific effects.

**Fig. 3.**
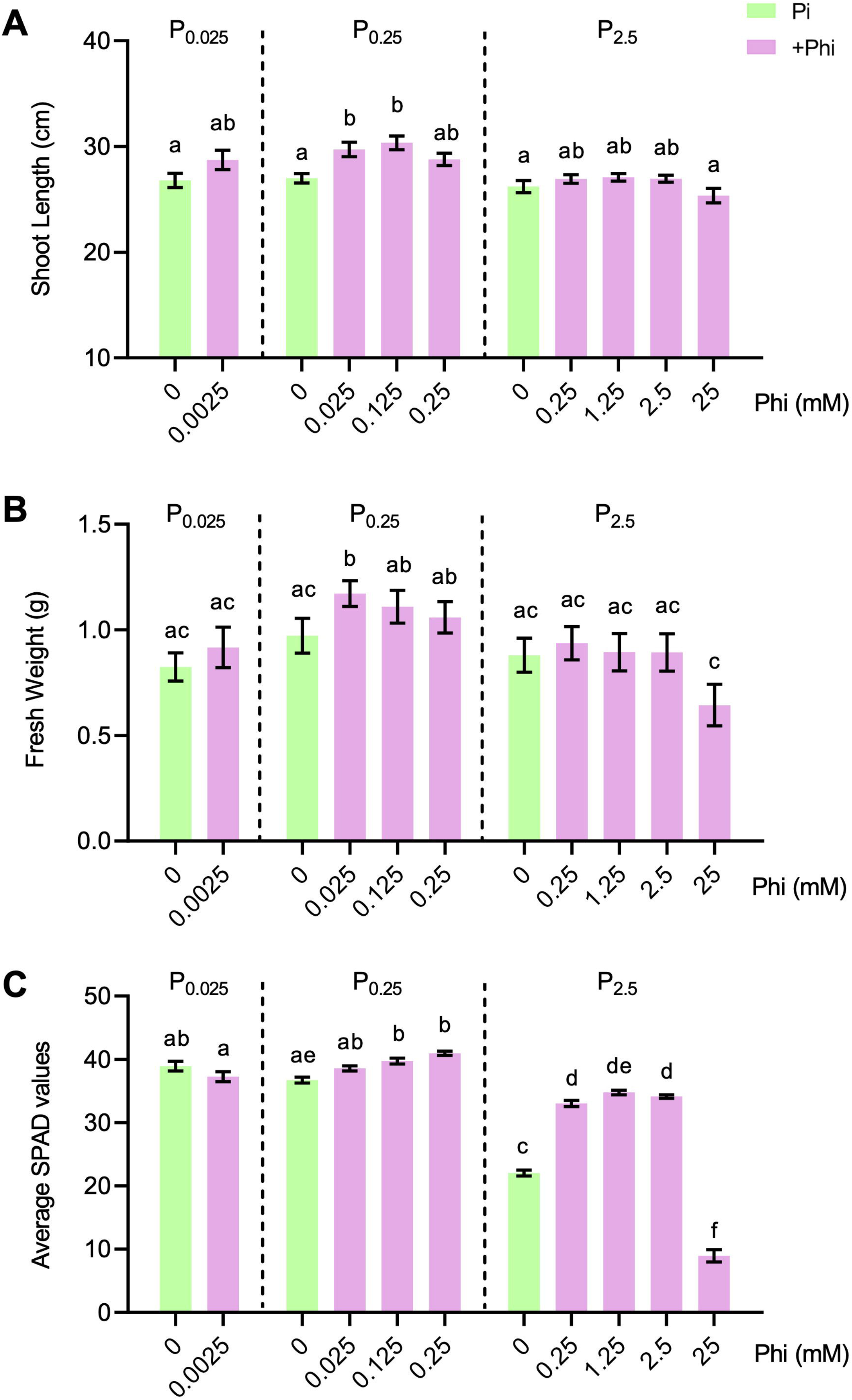
Phosphite promotes rice growth and increases photosynthetic pigment content in rice. Plants (*Oryza sativa* cv. Nipponbare) were pregerminated (1 week on water) and then grown under different Pi and Pi+Phi conditions for two weeks. Bar graphs represent mean values with error bars ± SEM from at least 8-10 biological replicates per treatment group. Letters above each bar represents statistical significance as per two-way ANOVA and Tukey’s multiple comparison tests, within and between the Pi and Pi+Phi treatments. Same letters indicate no statistical difference. (**A**) Average shoot length (cm) of rice plants. (**B**) Shoot fresh weight of rice plants. (**C**) Chlorophyll content as determined by SPAD measurements.

### Phosphite enhances resistance against *P. cucumerina* in Arabidopsis

Previously, we reported that Arabidopsis plants grown under a high Pi regime exhibited resistance to *P. cucumerina*, whereas treatment with low Pi enhanced susceptibility to this pathogen (Val-Torregrosa *et al*., 2022*b*). In this work, we investigated whether Phi supplementation influences the immune phenotype to *P. cucumerina* infection in Arabidopsis. For this, Arabidopsis plants grown with either Pi alone (P_0.025,_ P_0.25_, and P_2.5_) or in combination with Phi at a one tenth molar ratio to Pi were inoculated with a suspension of fungal spores. Overall, Phi application enhanced resistance to *P. cucumerina* infection (**Fig. 4A**). Quantitatively, fungal biomass was reduced approximately two to five-fold to the corresponding non-Phi treatment (**Fig. 4B**, pink bars). However, Phi application to P_0.025_ Arabidopsis plants had a detrimental effect, as revealed by extensive leaf death (**Fig. 4A**, lower panel). The observed reduction in fungal biomass that is observed in Phi-treated P_0.025_ plants (**Fig. 4B**) likely occurred at the cost of phytotoxicity.

**Fig. 4.**
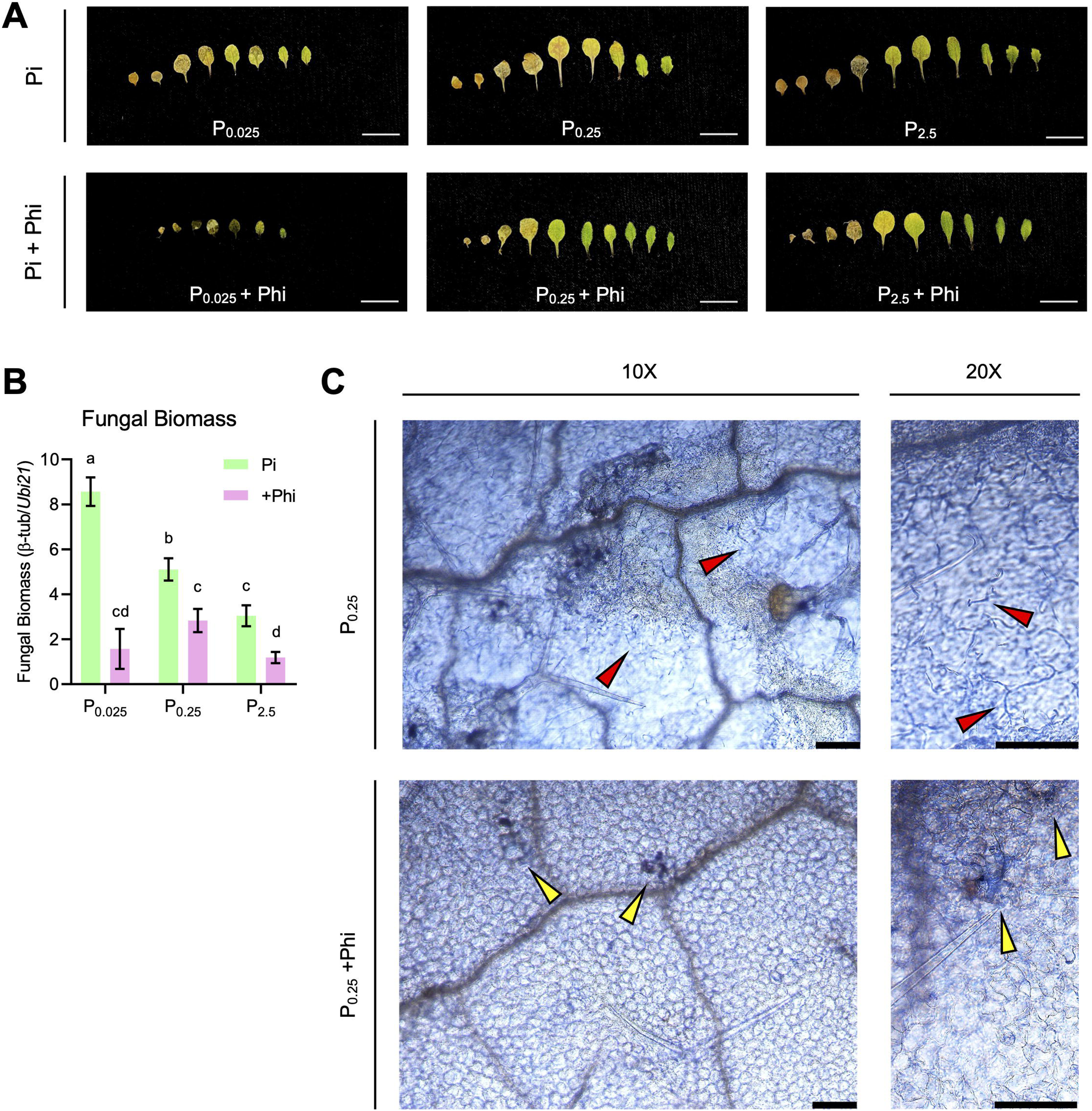
Protective effect of phosphite (Phi) supplementation in *Arabidopsis* against *P. cucumerina* infection. Plants were grown under 0.025 mM, 0.25 mM Pi or 2.5 mM Pi conditions, with or without Phi supplementation, and then spray-inoculated with *P. cucumerina* spores (5 x 10^5^ spores/mL). Plants were examined at 7 dpi. Three independent experiments were carried out with similar results. (**A**) Resistance to *P. cucumerina* infection in Phi-treated *Arabidopsis* plants. Upper panels, leaves of *P. cucumerina*-infected plants grown at the indicated Pi concentrations (Pi alone). Lower panels, leaves of *P. cucumerina*-infected Pi+Phi grown plants. Scale bars = 10mm. (**B**) Left panel, quantification of fungal biomass by qPCR using specific primers for *P. cucumerina* β*-tubulin* normalized against *Arabidopsis Ubiquitin21.* Letters above each bar represents statistical significance as per two-way ANOVA and Tukey’s multiple comparison tests, within and between the Pi and Pi+Phi treatments. (**C**) Trypan blue stained *P. cucumerina* infected leaves of plants grown under 0.25 mM Pi (P_0.25_), or with a combination of Pi and Phi (P_0.25_ + Phi_0.0025_). Red and yellow arrowheads indicate fungal hyphae and dead cell spots, respectively. 10X and 20X magnifications are indicated above panels. Scale bars = 100µm.

Trypan blue staining was used to visualize fungal structures and plants’ dead cells in the infected leaves of the above Arabidopsis plants. In untreated P_0.25_ plants, extensive fungal hyphal development was observed in leaves, indicating successful colonization (**Fig. 4C**, red arrowheads). In contrast, fungal colonization was absent in the co-treatment of P_0.25_ with Phi, though discrete patches of dead cells were visualized (**Fig. 4C**, yellow arrowheads). This pattern of scattered groups of dead cells is reminiscent of a hypersensitive response (HR), a form of localized cell death often associated with plant resistance to pathogen infection.

From these results, it can be concluded that supplementing Arabidopsis with Phi enhances resistance to *P. cucumerina* in Arabidopsis in a Pi-dependent manner (e.g., under optimal and high Pi supply). Under low Pi supply, however, Phi has detrimental effects in Arabidopsis plants. Knowing that Phi cannot be metabolized by the plant, Phi supplementation to Pi-deficient Arabidopsis plants might well prevent Pi from fulfilling its essential biochemical functions, thus, negatively affecting normal metabolic processes in the plant. These findings together with those presented above on the effect of Phi on plant growth support that, under adequate Pi levels, Phi supplementation can improve performance and disease resistance in Arabidopsis.

### Phosphite reverses Pi-induced susceptibility to rice blast and bakanae disease

We investigated the capability of Phi treatment to suppress important fungal diseases in rice, namely blast and bakanae diseases. *M. oryzae* is a foliar pathogen and the causal agent of blast disease not only in rice but also in other cereals like wheat, barley and millet (Wilson and Talbot, 2009; Wang and Valent, 2017). Indeed, blast is one of the most destructive diseases of cultivated rice and represents a significant threat to rice production worldwide (Fernandez and Orth, 2018). The bakanae disease (or ‘foolish seedling’ disease) caused by the seed- and soil-borne pathogen *F. fujikuroi*, is also a widespread disease in major rice-growing countries (An *et al*., 2023). Root colonization by *F. fujikuroi* in rice has been previously described (Chen *et al*., 2020; Aragona *et al*., 2021). Importantly, treatment with high Pi has been shown to enhance susceptibility to infection by *M. oryzae* and *F. fujikuroi*, also known as Phosphate-Induced Susceptibility (Campos-Soriano *et al*., 2020; Martín-Cardoso *et al*., 2025).

Rice plants were grown under different Pi regimes, 0.025mM Pi, 0.25mM Pi, and 2.5mM Pi, either alone or in combination with Phi. Then, rice plants were spray-inoculated with a spore suspension of *M. oryzae* (Guy11 strain), and disease progression was evaluated over time. Under P_0.025_, rice plants exhibited inherent resistance to blast infection, as revealed by visual inspection of disease symptoms, lesion area quantification and measurement of fungal biomass (**Fig. 5A-C**). In contrast, growing rice plants under a high Pi regime (e.g., P_2.5_ plants, without Phi supplementation) results in susceptibility to infection by the rice blast fungus (**Fig. 5A-C**), which is consistent with results previously reported (Campos-Soriano *et al*., 2020). Notably, Phi supplementation to rice plants grown under optimal or high Pi regimes increased blast resistance, whereas Phi supplementation to Pi-deficient plants (P_0.025_ plants) did not significantly alter the immune phenotype (**Fig. 5A-C**).

**Fig. 5.**
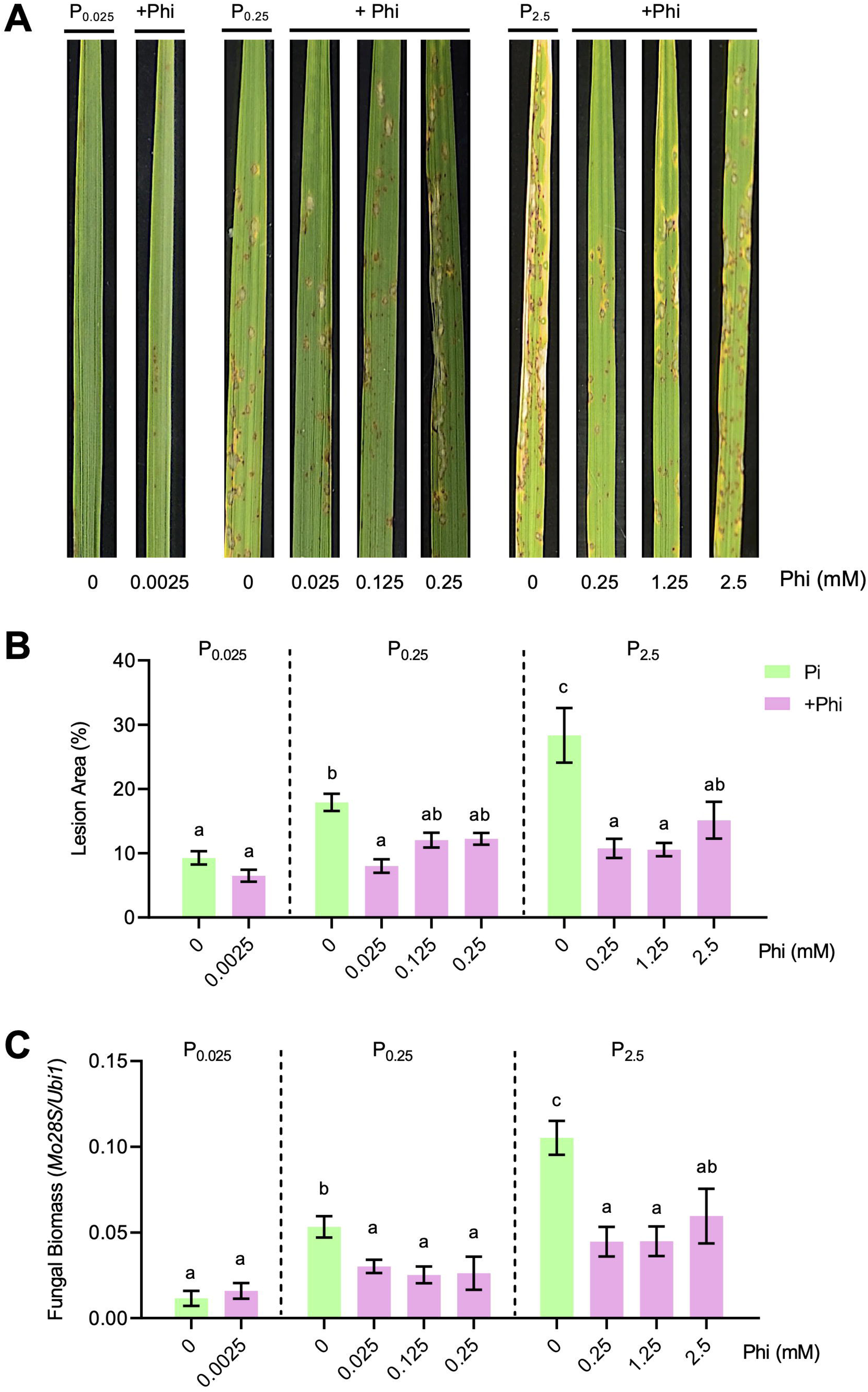
Phosphite reverts Pi-induced blast susceptibility of rice plants. (**A**) Representative images of 3-4 leaf stage rice (cv. Nipponbare) plants leaves grown for 2 weeks under 0.025mM Pi, 0.25mM Pi, or 2.5mM Pi condition, without Phi supplementation or with Phi at the indicated concentrations. Images correspond to the 3^rd^ leaf of plants, 7 days post infection. (**B**) Quantification of blast lesion area as a percentage of total leaf area across Pi and Pi+Phi treatments. (**C**) Fungal biomass was quantified by qPCR using specific primers for *M. oryzae 28S* gene normalized to rice *Ubi1* gene. Bars plots in B and C represent mean values ± SEM of 4 biological replicates (6-8 plants per replicate). Letters above each bar indicate statistical significance based on two-way ANOVA followed by Tukey’s multiple comparison test, comparing values within and between Pi and Pi+Phi treatments. Bars sharing the same letter are not significantly different from each other.

In agriculture, Phi can be used in the form of potassium phosphite (KPhi, KH_2_PO_3_) or sodium phosphite (NaPhi, Na_2_HPO_3_·5H_2_O). In the experiments so far, KPhi was used in both Arabidopsis and rice. When comparing the effect or one or another Phi salt on disease resistance in rice, the level of disease suppression was comparable between KPhi and NaPhi, indicating that both salts confer protection against rice blast (Supplementary Fig. S3).

The effect of Phi in protecting rice plants against the blast fungus was examined in several commercial rice varieties of *japonica* that are cultivated in temperate regions, namely Argila, Baixet, Bomba, and Guara. Plants were grown under high Pi supply (2.5mM Pi, a condition that enhances blast susceptibility in Nipponbare plants), with or without Phi (at a 1/10 Pi/Phi ratio). Phi application conferred improved resistance in all four rice varieties, evidenced by visual inspection of *M. oryzae* infected leaves, and further validated by quantification of lesion area and fungal biomass which were reduced in Phi-supplemented plants (Supplementary Fig. S4A, B). However, the protective effect of Pi was found to vary depending on the rice genotype (e.g., Argila benefited the most from Phi treatment during *M. oryzae* infection). SPAD measurements revealed higher chlorophyll content in plants supplied with Phi in all the rice varieties, thus, reflecting improved health under Phi treatment (Supplementary Fig. S4C). These results highlight the potential of Phi to attenuate Pi-induced blast susceptibility in commercial rice varieties, hence, for management of rice blast, at least in Argila, Baixet, Bomba and Guara plants.

Knowing that treatment with high Pi increases susceptibility to pathogen infection, and that Phi application suppresses Pi-induced susceptibility in rice plants, it was of interest to assess whether Phi application alters Pi content in rice tissues. In Phi-treated P_0.25_ and P_2.5_ plants, Pi content was consistently lower than in Pi-only controls in both leaf and root tissues (Supplementary Fig. S5A, B). The same trend was observed in Nipponbare and the cultivated rice varieties Argila, Baixet, Bomba and Guara assessed here (Supplementary Fig. S5B).

In this work, we also assessed whether Phi supplementation can counteract the negative effect of Pi on Bakanae disease. For this, rice plants were grown under different Pi regimes, with or without Phi, and roots were subsequently inoculated with *F. fujikuroi* spores. As expected, the roots of infected plants were shorter and showed necrotic regions compared to Pi only controls (**Fig. 6A**). Remarkably, Phi supplementation to plants grown under optimal and high Pi conditions (P_0.25_ + Phi, P_2.5_ + Phi) was accompanied by a reduction in fungal biomass and a lower Disease Severity Index relative to their Pi-only counterparts (**Fig. 6B, C**), confirming higher resistance in plants supplemented with Phi. These observations support that Phi supplementation enhances resistance to Bakanae disease in rice under adequate or high Pi accumulating plants.

**Fig. 6.**
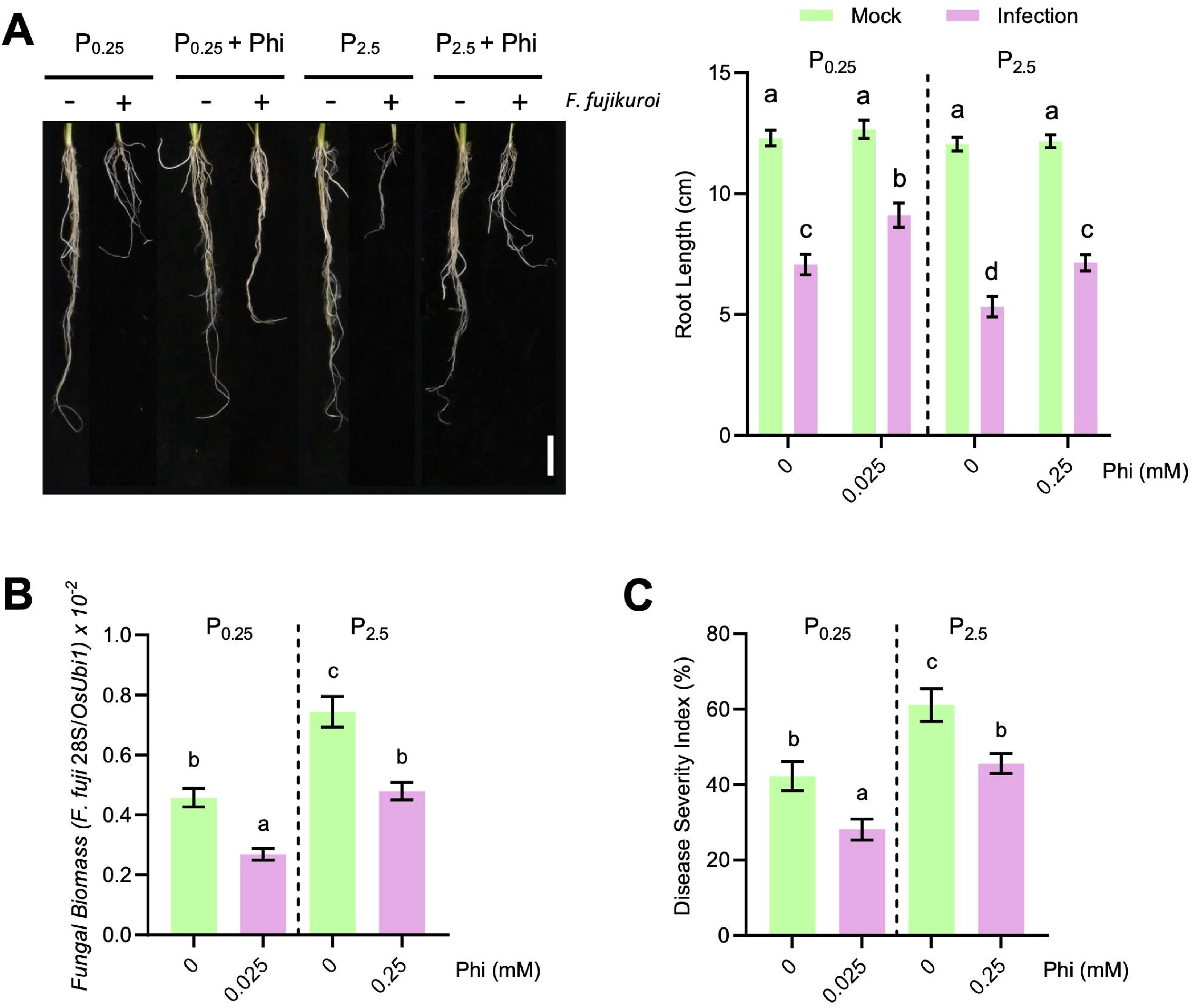
The application of Phi influences root growth and enhances resistance to bakanae disease. (**A**) Left panel, representative images of roots of rice seedlings grown under different Pi regimes (0.025 mM, 0.25 mM, and 2.5 mM Pi) with or without Phi supplementation (1/10th Phi:Pi ratio) and either mock-inoculated (-) or inoculated with *Fusarium fujikuroi* (+). Scale bar = 1 cm. Right panel, the corresponding bar graph shows root length measurements of mock and infected plants under each treatment. (**B**) Quantification of *F. fujikuroi* fungal biomass by qPCR using specific primers for fungal 28S rDNA and normalized to rice *Ubiquitin1* gene expression. (**C**) Disease severity index (DSI) expressed as percentage of infected seedlings showing bakanae symptoms. Bars represent mean values ± SEM from four biological replicates (6–8 plants per replicate). Letters above bars indicate statistical significance based on two-way ANOVA followed by Tukey’s multiple comparison test. Bars sharing the same letter are not significantly different (p < 0.05).

Collectively, results from infection experiments demonstrated that Phi supplementation reverses Pi-induced susceptibility to blast and bakanae disease. Phi supply does not affect the resistance already present under low Pi-deficient conditions. The protective effect of Phi against *M. oryzae* occurs in different *japonica* rice varieties, such as Nipponbare (model *japonica* variety for studies on functional genomics), and other temperate *japonica* rice varieties, such as Argila, Baixet, Bomba and Guara.

### Foliar application of phosphite enhances blast resistance

Results shown above demonstrated the efficacy of Phi in protection against the rice blast fungus when Phi is applied to the soil through fertigation. In this work, we also investigated whether foliar application of Phi serves to protect rice plants from *M. oryzae* infection. For this, rice plants at the 3-4 leaf stage grown under P_0.25_ were sprayed with Phi solution and, 24 hours later, the rice plants were spray-inoculated with *M. oryzae*. As shown in **Fig. 7A**, foliar application of Phi significantly reduced blast symptoms compared to untreated plants. A visible reduction in disease severity was observed, which was supported by quantitative analysis of diseased leaf area showing a ∼60% decrease in lesion area (*P* < 0.01, Student’s *t*-test) (**Fig. 7B**). Fungal biomass was also significantly reduced in Phi-sprayed plants (*P* < 0.05, Student’s *t*-test) (**Fig. 7C**), confirming improved resistance to blast upon Phi treatment. These findings highlight the effectiveness of foliar Phi application in enhancing resistance to *M. oryzae*.

**Fig. 7.**
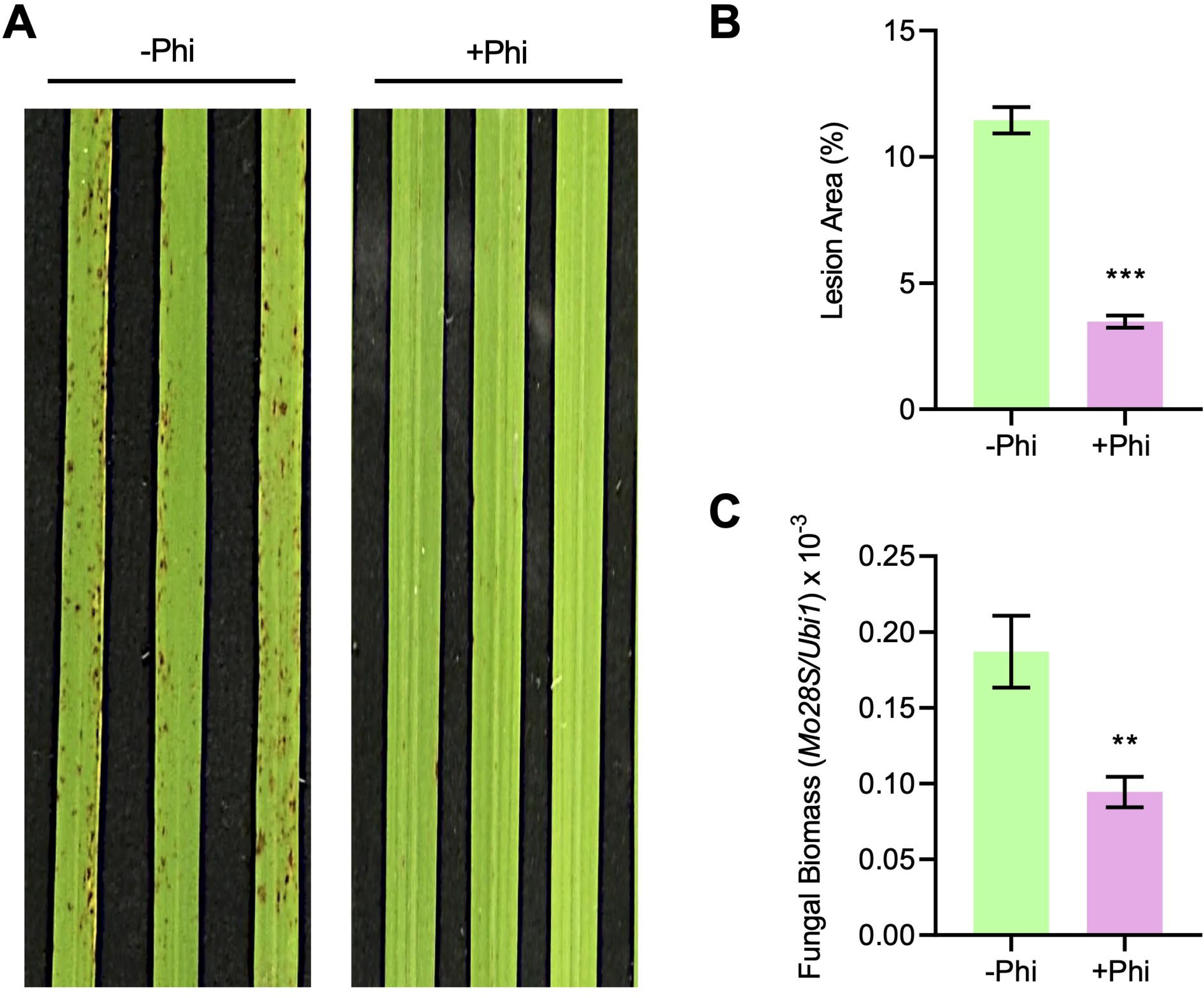
Foliar application of Phi enhances blast resistance. Plants at the 3-4 leaf developmental stage, grown under 0.25 mM Pi supply, were sprayed with a solution of 10 mM KPhi (+Phi). Control plants were sprayed with sterile water (-Phi). Infection with *M. oryzae* was carried out 24 hours later, and disease symptoms were evaluated at 7 dpi. (**A**) Representative images of disease symptoms (3^rd^ leaf of plants at the 3-4 leaf stage). (**B**) Percentage of diseased leaf area. (**C**) Fungal biomass quantification by quantitative PCR analysis using specific primers for *Mo28S* and normalized to rice *Ubi1* gene (right panel). Bars represent ± SEM of 4 biological replicates, 6 plants per replicate and statistical significance tested by Student’s *t*-test, with \**p* < 0.05, \*\**p* < 0.01.

### Transcriptional reprogramming in rice leaves in response to phosphite treatment

To get molecular insight into the potential of Phi to revert the adversities of high Pi treatment, a transcriptome deep sequencing was performed on rice leaves from P_2.5_ plants supplemented or not with Phi. 1,546 transcripts were found differentially expressed due to Phi (adjusted *P* ≤ 0.05, Wald’s test), of which 888 genes were up-regulated and 658 genes were down-regulated (Supplementary Table S2). Up-regulated transcripts were not only more abundant but also displayed the highest fold-changes (**Fig. 8A**), suggesting that Phi supply mainly imposed activation of transcriptome responses. However, functional annotation of changing transcripts based on Gene Ontology (GO) terms (adjusted *P* < 0.05, Fisher’s Exact test) revealed that, despite being up-regulated transcripts more prevalent, they represented fewer biological processes than the down-regulated (16 vs 65 non-redundant terms) (**Fig. 8**; Supplemental Table S2). Thus, the transcriptome reprogramming under Phi might include a higher level of orchestration to efficiently repress pathways.

**Fig. 8.**
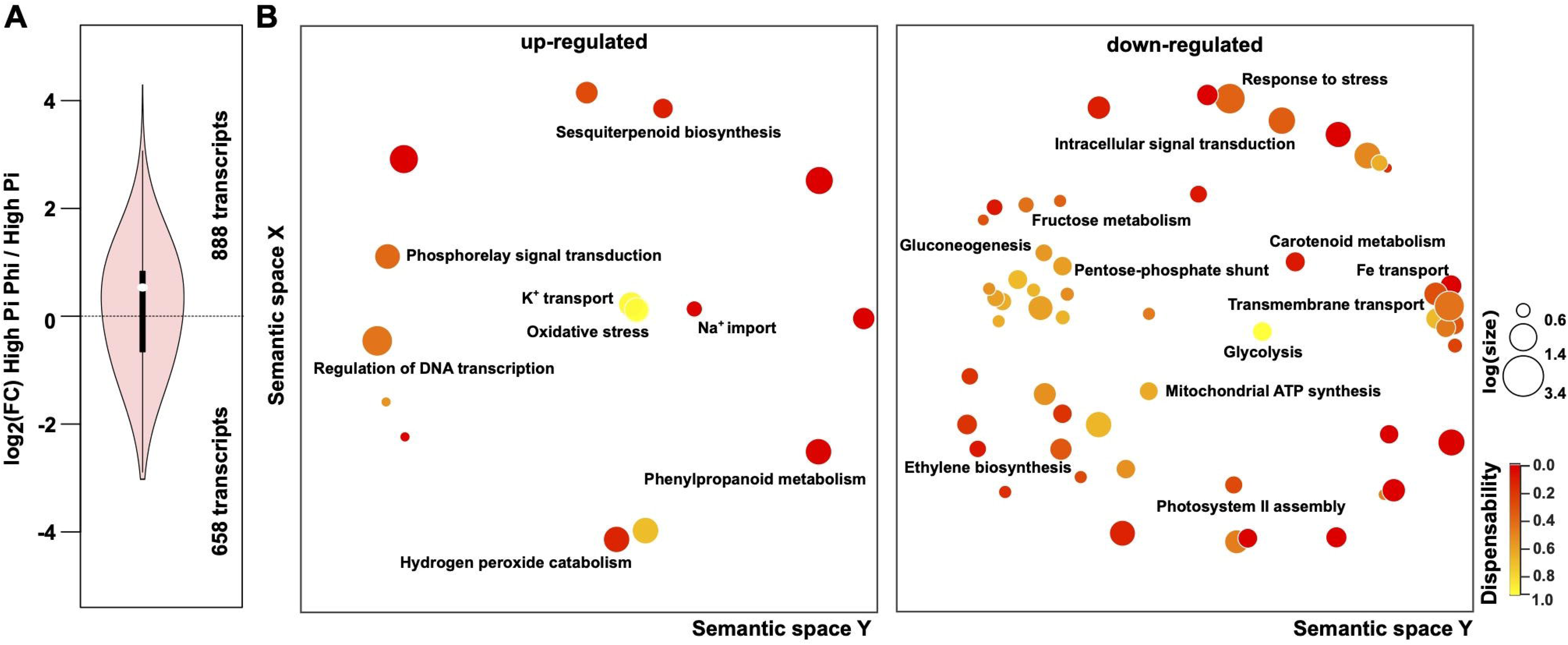
Transcriptome profiling of plants co-treated with Phi and high Pi. RNA-Seq data from leaves of rice plants cultivated under 2.5 mM Pi were supplemented or not with 0.25 mM Phi were filtered for transcripts significantly changing (adj *P* ≤ 0.05, Wald’s test). (**A**) Violin plot depicting the distribution of the log_2_ fold-change (FC). (**B**) Bubble plots summarizing the Gene Ontology terms significantly enriched (adj *P* ≤ 0.05, Fisher’s exact test) for changing transcripts according to the sign (up-regulated or down-regulated). Indicated are the most relevant terms.

As depicted in **Fig. 8B**, the addition of Phi to P_2.5_ plants promoted terms mainly related to the sensing and transduction of signals (“Regulation of DNA-templated Transcription”, “Phosphorelay Signal Transduction System”), oxidative stress responses (“Cellular Responses to Oxidative Stress”, “Response to Hormones”, “Metabolic Process”) along with transport of ions (“Potassium and Sodium Transmembrane Transport”) (**Fig. 8**; Supplementary Table S3). Conversely, down-regulated transcripts significantly enriched processes related to central metabolism of carbon (“Glycolysis/Gluconeogenesis”, “Pentose-phosphate shunt”), stress-responsive mechanisms (“Ethylene Biosynthetic Process”, “Response to Stress”, “Chloroplast Organization/Photosystem II assembly”, “Carotenoid Metabolic Processes”) and signaling (“Intracellular Signal Transduction”) (**Fig. 8**; Supplementary Table S4). Among DEGs, genes related to proton motive force-driven mitochondrial ATP synthesis (Supplementary Table S5), as well as genes involved in Phosphorelay Signal Transduction (Supplementary Table S6) were also found to be regulated by Phi. In consequence, Phi supply on rice plants treated with P_2.5_ imposed a less constraint state that circumvented the necessity to transcriptionally force metabolic pathways and stress-responsive mechanisms, including sustaining signalling pathways depending on phosphorylation, under high Pi.

## Discussion

In agriculture, Phi has multiple applications including protection against microbial pathogens and weeds, and also serves as a biostimulant (Li *et al*., 2025). Phi has been particularly useful in managing diseases caused by oomycetes such as *Phytophthora* spp. and *Hyaloperonospora arabidopsidis* (Massoud *et al*., 2012; Achary *et al*., 2017; Havlin and Schlegel, 2021; Mohammadi *et al*., 2021; Li *et al*., 2025). As previously mentioned, two possible mechanisms have been proposed to explain Phi-mediated protective effects from diseases consisting of direct inhibition of fungal growth and, indirectly through the induction of the plant’s own defense mechanisms (Achary *et al*., 2017; Mohammadi *et al*., 2021; Trejo-Téllez *et al*., 2024). At present, however, the exact mechanisms by which Phi protects crop species from pathogen infection remain poorly understood.

In this study, we show that Phi treatment has a differential effect on the growth of phytopathogenic fungi. Distinct effects of Phi in fungal growth might be related to differences in the integration of Pi homeostasis with fungal metabolic processes (e.g., carbon metabolism, ATP and nucleic acid biosynthesis), signaling pathways, or membrane integrity (Bhalla *et al*., 2022). Further, Pi serves as structural and metabolic molecules in fungi, and its availability would then control fungal growth and development (Tan *et al*., 2025). Phi might target specific fungal processes affecting fungal growth, which might differ among pathogens. As a consequence, although Phi cannot be metabolized, Phi would interfere with normal fungal growth and development.

We also show that the application of Phi exerts a dual action in Arabidopsis and rice plants. Thus, Phi has been shown to influence Arabidopsis and rice growth in a differential and concentration-dependent manner. Not only the plant species, but also the plant Pi status and the Pi/Phi relative concentrations are important factors in determining the effect of Phi treatment on plant growth. For instance, while the application of Phi has a biostimulant effect in Arabidopsis plants under sufficient Pi, it negatively affects Arabidopsis growth under Pi deficiency. As Phi cannot replace Pi for direct P nutrition, these findings suggest that Phi might aggravate stress caused by Pi deficiency in Arabidopsis, likely, interfering with Pi starvation responses in the plant. Inconsistent findings are, however, found in the literature on the effect of Phi on Pi starvation responses in plants. In some cases, Phi suppresses plant responses that would normally be activated under Pi deprivation in Arabidopsis, *Brassica napus* or tomato plants, thus, hindering plant survival under Pi limiting conditions (Carswell *et al*., 1996; Ticconi *et al*., 2001; Varadarajan *et al*., 2002). Arguing against Phi-mediated suppression of Pi starvation responses, Phi has been shown to induce the expression of Pi transporters and phosphatase genes in alfalfa, a typical response to Pi starvation (Li *et al*., 2022). Our findings indicate that, under Pi limiting conditions, Phi supplementation leads to inhibition of Arabidopsis growth, whereas the addition of Phi to rice plants under low Pi supply does not penalize plant growth. Different results can be also found in the literature related to the effect of Phi on the plant PSR by either attenuating or activating this response which might well be related to differences in the experimental conditions used for Pi and Phi treatment. Another finding of this study relates to the Pi/Phi ratio used for treatment of plants, an aspect that should be carefully considered in plants grown under Pi sufficiency or high Pi conditions. Optimal effects of Phi treatment were observed at a one tenth Phi to Pi ratio. Clearly, a combination of different factors might account for complexity of results across different studies, including species’ unique characteristics, and differences in growth conditions (e.g., Pi and Phi supply) or Pi/Phi ratios. Under this scenario, it can be concluded that Phi application can be beneficial to the plant when Pi and Phi are adequately supplied. An adequate Pi nutrition is essential to avoid potential deleterious effects of Phi on plants, an information that might help in delineating conditions benefiting the most the rice plants from Phi application.

A central theme of this study was to decipher the impact of Phi application on disease resistance in Arabidopsis and rice plants as models for dicot and monocot studies. In Arabidopsis, Phi treatment increases resistance to *P. cucumerina* infection in plants grown under sufficient or high Pi regimes. Resistance to *P. cucumerina* in Phi-treated Arabidopsis plants is accompanied by localized cell death. This response is reminiscent of a hypersensitive response (HR), a plant’s natural defense system that restricts pathogen colonization and confers disease resistance in Arabidopsis. In another *Brassica* species, *B. napus*, Phi was also found to accelerate programmed cell dead in suspension cells under Pi deficiency (Singh *et al*., 2003).

Importantly, results here presented demonstrated that Phi application to rice plants suppresses Pi-induced susceptibility to blast and bakanae disease. The protective effect of Phi against these fungal diseases was observed in the cultivar Nipponbare (widely used in functional genomic research), as well as in commercial rice varieties (e.g., Argila, Baixet, Bomba, Guara). Both potassium and sodium salts of Phi, were found to enhance blast resistance. Furthermore, both soil application and foliar application of Phi through spray provided substantial protection against blast disease, these results aligning with those reported in other crops on pathogen resistance conferred by foliar application of Phi (Vinas *et al*., 2020; Mehta *et al*., 2022). Hence, Phi application to rice plants, either soil application or foliar application, appears to be a valuable management strategy to fight *M. oryzae* infection in rice cultivation, being the foliar application a more efficient solution in terms of timming.

From the results here presented, several possibilities can be envisaged to explain the protective effect of Phi against pathogen infection in Arabidopsis and rice plants that involves an effect of Phi in one or another partner, plant and pathogen. In Arabidopsis, this study revealed that Phi exerts a direct inhibitory effect on *P. cucumerina* growth, whereas earlier studies reported that Phi treatment activates Arabidopsis defense responses (Pérez-Zavala *et al*., 2024). The observed resistance to *P. cucumerina* in Phi-treated Arabidopsis plants might well result from a combination of direct inhibition of fungal growth (present work) and activation of plant defense responses (Pérez-Zavala *et al*., 2024). Then, in the Arabidopsis/*P. cucumerina* pathosystem, Phi would exert its protective effects by acting on the two interacting partners, the host plant and the fungus.

Conversely, *M. oryzae* growth was not inhibited effectively by Phi. Thus, based on differences observed in the response to Phi supplementation in the fungal pathogens here investigated, an inhibitory effect on growth of phytopathogens by Phi cannot be generalized. Here, it is worthy to mention that Pi is not only an essential nutrient for fungal growth but also participates in signal transduction and adaptation to environmental variations (Tan *et al*., 2025). It can be reasoned that Phi might negatively affect fungal growth (e.g., *P. cucumerina*, *F. fujikuroi* growth) by competitively disrupting Pi-dependent metabolic pathways in fungal cells, including phosphorylation reactions (Achary *et al*., 2017; Havlin and Schlegel, 2021). On the other hand, Pi accumulation in rice leaves has been shown to promote the expression of *M. oryzae* effectors, hence, stimulating fungal pathogenicity (Martín-Cardoso *et al*., 2024). Phi might well compete with Pi for the production of *M. oryzae* effectors, hence, attenuating Pi-mediated stimulation of *M. oryzae* effectors production. The possibility that Phi exposure provokes alterations in Pi metabolism and/or *M. oryzae* pathogenicity should be considered. In other studies, it was demonstrated that Phi interferes with fungal metabolism, either through inhibition of key phosphorylation reactions or by altering nucleotide pools (Achary *et al*., 2017; Dempsey *et al*., 2018). Clearly, further investigation is required to clarify Phi’s effect on fungal growth and pathogenicity.

In addition to a direct effect of Phi on fungal growth, alternative—though not mutually exclusive—mechanisms acting on the host plant likely contribute to the enhanced disease resistance observed. In this direction, our transcriptomic profiling revealed that, under high Pi conditions, Phi co-treatment promoted a more resilient physiological state. Notably, Phi alleviated the activation of general stress-related pathways and dampened the stress-induced upregulation of central carbon metabolism genes (Fig. 8), suggesting a restoration of metabolic homeostasis. In parallel, Phi supply induced the accumulation of transcripts associated with oxidative stress mitigation. Together, these responses could suggest that Phi exerts a protective effect, reduces the need for costly stress-driven metabolic reprogramming under high Pi, thereby preserving resources for growth and enhancing the plant’s capacity to respond to subsequent environmental challenges.

This study provides mechanistic insights into molecular processes that are specifically regulated by Phi in high-Pi rice plants. Specifically, this study demonstrated that Phi treatment has a strong impact on the expression of genes involved in Phosphorelay signal transduction. It is well known that plants evolved a phosphorelay signal transduction system that involves a multi-step series of phosphorylation events that provide a robust framework of plants to sense and respond to environmental stimuli (Pekárová *et al*., 2016; Sharan *et al*., 2017). In Arabidopsis, this phosphorelay signaling mechanism involves different types of proteins, namely a sensory histidine kinase, a phosphotransfer protein, and a response regulator. Phi treatment appears to modulate the expression of Histidine Kinase 3 (HK3, the primary sensor protein) and several response regulators in the phosphorelay signal transduction system. In other studies, this complex system was reported to regulate salt sensitivity and resistance against bacterial and fungal pathogens in Arabidopsis (Pham *et al*., 2012). Thus, results here presented underscore the importance of Phi in regulating phosphorylation steps involved in signaling mechanisms that allow the plant to sense and respond to environmental cues (i.e., Pi stress). This regulation, in turn, would contribute to counteract the negative effect that Pi accumulation has on disease resistance.

Apart from the regulation of genes involved in phosphorelay signal transduction, our transcriptome analysis of Phi-treated rice plants revealed Phi-triggered alterations in genes involved in ethylene signaling in the absence of pathogen infection. Ethylene biosynthesis and signaling has been reported to be required for basal resistance against the blast fungus *M. oryzae* in rice (Helliwell *et al*., 2016). Phi-mediated regulation of genes associated with ethylene signaling might well contribute to revert Pi-induced blast susceptibility in Phi-treated rice plants. In Arabidopsis, Phi treatment has been shown to modulate the expression of genes associated with hormonal regulation of immune responses, such as ABA, SA and JA/ET pathways (Eshragi *et al*., 2011; Sánchez-Vallet *et al*., 2012; Pérez-Zavala *et al*., 2024). Although increasing evidence support a link between Phi and hormone signaling in plant defense, further studies are fundamental to understand how Phi regulates these processes in rice.

Together, results presented in this work provide evidence that Phi functions as a molecule capable of reverting Pi-induced blast susceptibility in rice by regulating the expression of genes involved in phosphorylation signaling and ethylene signaling. We propose that Phi treatment makes the rice plant to be in a primed status for attenuation of Pi-induced blast susceptibility. Phi is already commercially and used in agriculture as a biostimulant and antimicrobial compound. From the results presented, it can be concluded that gaining benefits from the application of Phi is highly dependent not only on the basal level of Pi in the plant but also on the Pi to Phi ratio applied. Thus, the efficiency of its use in agriculture needs to be evaluated in the context of current Pi-based fertilization regimes on a case-by-case basis. Phi’s effectiveness in protecting rice against *M. oryzae* and *F. fujikuroi* is particularly important, as Phi can suppress Pi-induced susceptibility to blast and bakanae diseases, two devastating fungal diseases in rice that significantly threaten rice production. As overuse of Pi fertilizers and pesticides continues to pose ecological challenges, Phi’s capability to improve disease resistance will contribute to maintain sustainable rice production.

## Abbreviations

(ABA): Abscisic acid
(CDPKs): calcium-dependent protein kinases
(DEGs): differentially expressed genes
(ET): ethylene
(JA): jasmonic acid
(MAPK): mitogen-activated protein kinases
(MS medium): Murashige–Skoog medium
(Pi): phosphate
(Phi): phosphite
(P): phosphorus
(SA): salicylic acid
(SPAD): Soil Plant Analysis Development.

## Supplementary data

The following supplementary data are available at JXB online.

**Fig. S1**. Schematic representation of phosphate (Pi) and phosphite (Phi) treatment combinations used in this study.

**Fig. S2.** Inhibition of *P. cucumerina* growth by Phi in solid medium.

**Fig. S3.** Effect of phosphite salts on rice blast resistance.

**Fig. S4.** Protective effect of Phi against *M. oryzae* infection in *japonica* rice cultivars grown under high Pi supply.

**Fig. S5.** Free Pi content in rice varieties supplemented, or not, with Phi.

**Table S1**. List of oligonucleotides used in this study.

**Table S2**. Differentially expressed genes due to the application of phosphite to high phosphate treatments.

**Table S3**. Functional annotation of up-regulated transcripts.

**Table S4**. Functional annotation of down-regulated transcripts

**Table S5**. List of transcripts related to proton motive force-driven mitochondrial ATP synthesis.

**Table S6**. List of transcripts related to phosphorelay signal transduction system.

## Acknowledgements

We thank Montse Amenós (Microscopy and Imaging Facility) and the Greenhouse facility at the CRAG for technical assistance.

## Authors contributions

BSS: designed the research; MDM: performed experiments, investigation and formal analysis; HM-C and GB contributed to *F. fujikuroi in vitro* growth and infection experiments; MA, contributed with *M. oryzae* infection in rice varieties; AG-M, contributed to RNASeq analysis (data curation, formal analysis and visualization); BSS, Funding acquisition and Project administration; MDM and BSS, Writing-original draft; HM-C and A.G-M, writing - review & editing. All authors read and approved the final manuscript.

## Data availability

The RNA-Seq data underlying this article are available in the Gene Expression Omnibus (GEO) Database (https://www.ncbi.nlm.nih.gov/geo/) and can be accessed with accession number GSE313364.

