## Supplementary Figures for "Phosphite, an analog of phosphate, counteracts Phosphate Induced Susceptibility of rice to the blast fungus *Magnaporthe oryzae*"

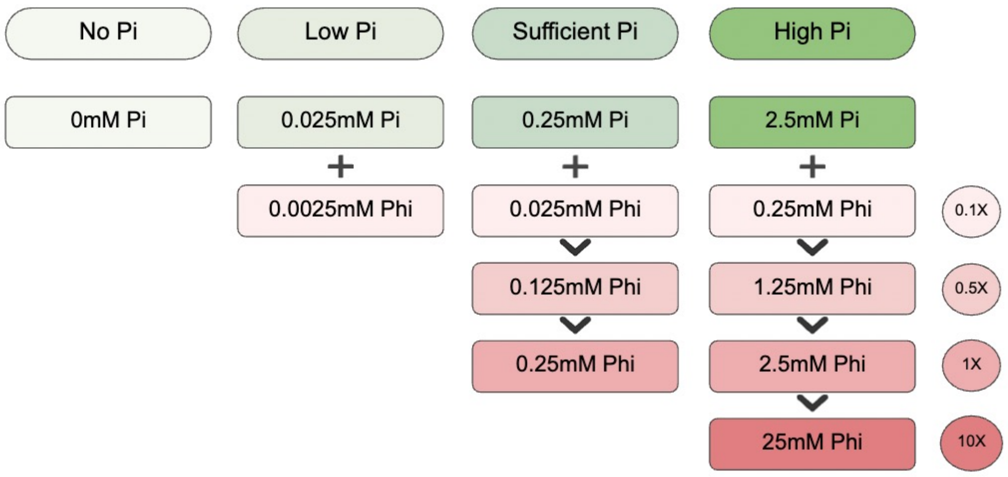


**Supplementary Figure S1**. Schematic representation of phosphate (Pi) and phosphite (Phi) treatment combinations used in this study. Four Pi conditions were tested: No Pi (0mM), 0.025mM Pi, 0.25mM Pi, and 2.5mM Pi, indicated in green boxes. To assess the concentration-dependent effects of Phi, a range of Phi:Pi ratios were tested, at 0.1X, 0.5X, 1X, and 10X relative to the corresponding Pi level in each condition. They included 0.0025mM, 0.025 mM, 0.125mM, 0.25mM, 1.25mM, 2.5mM, and 25mM Phi, indicated in pink boxes.

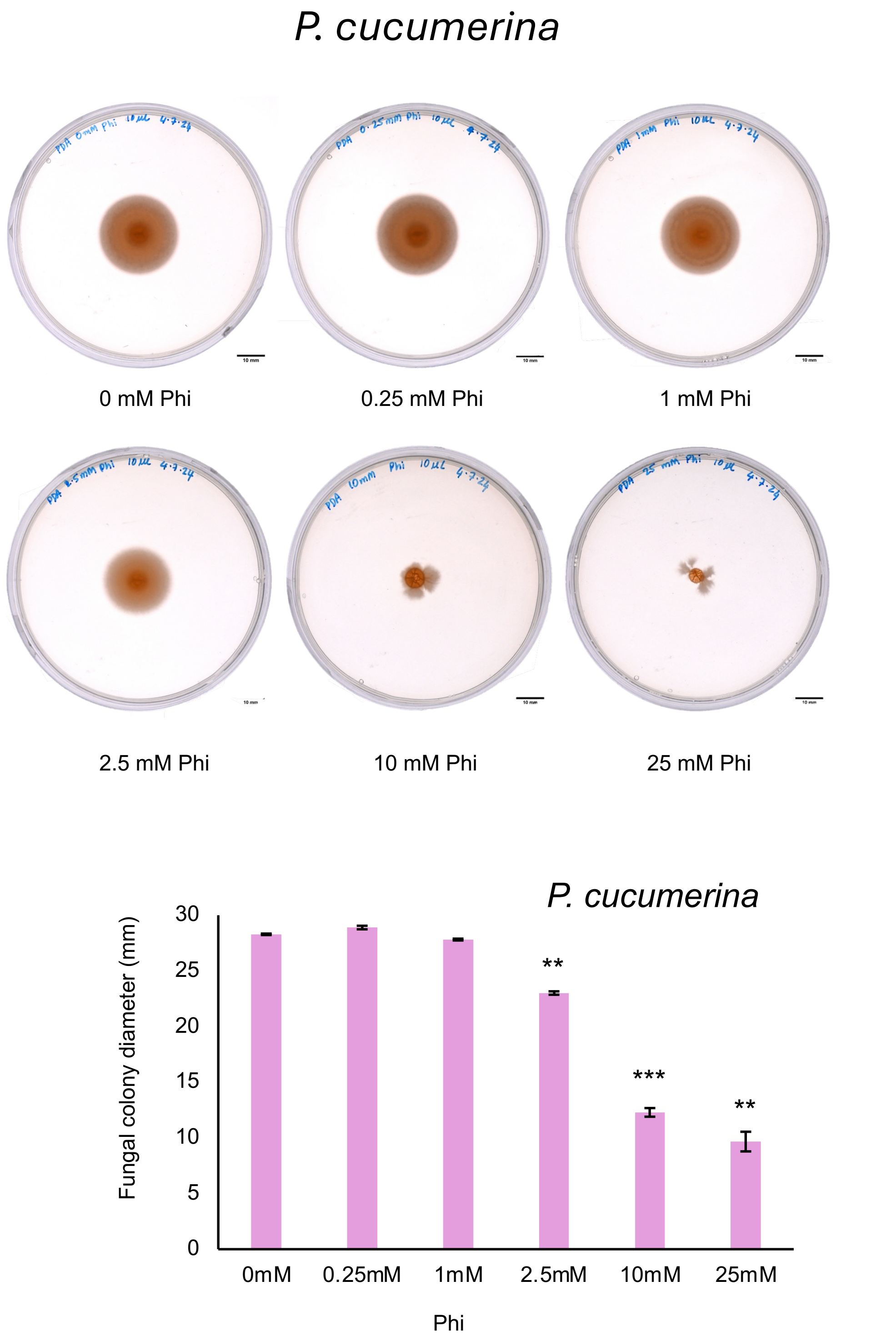


**Supplementary Figure S2.** Inhibition of *P. cucumerina* growth by Phi in solid medium. Colony morphology of *P. cucumerina* grown on PDA plates supplemented with increasing concentrations of Phi (0-25 mM). Scale bars = 10mm. Lower panel, fungal growth quantified by measuring colony diameter in mm. Statistical significance tested by Student's *t*-test with * $p<0.05$, ** $p<0.01$, *** $p<0.001$ and N=3 compared to preceding concentration.

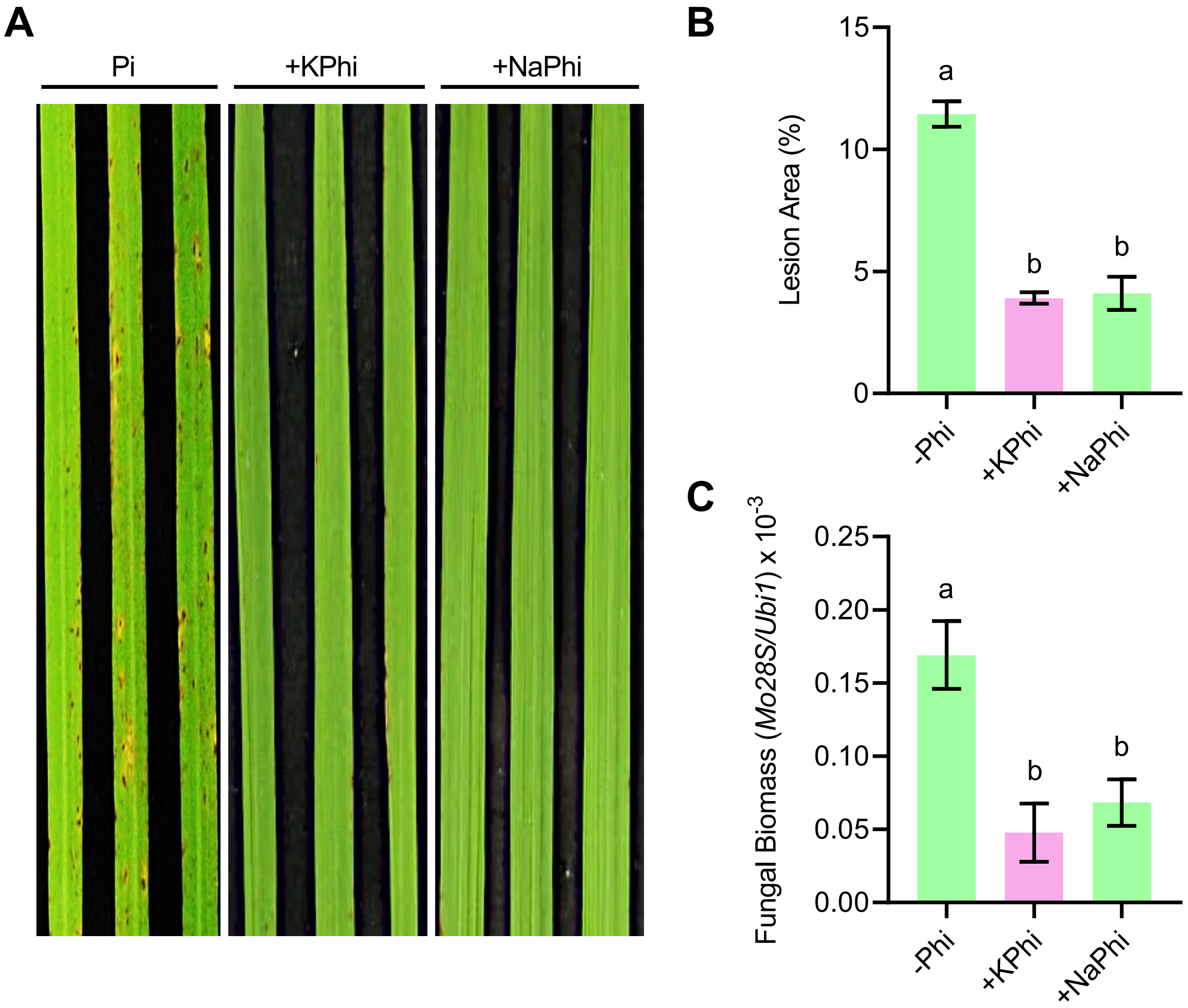


**Supplementary Figure S3.** Effect of phosphite salts on rice blast resistance. Rice (cv. Nipponbare) plants were grown under 0.25mM Pi supply (Pi) supplemented with either potassium phosphite (+KPhi, 0.025mM) or sodium phosphite (+NaPhi, 0.025mM). Plants were spray inoculated with a spore suspension of *M. oryzae*. (**A**) Representative images of disease symptoms at 7 days post-inoculation with *M. oryzae* spores. (**B**) Quantification of lesion area (3$^{rd}$ leaf of plants at the 3-4 leaf stage). (**C**) Quantification of fungal biomass by qPCR using specific primers of *M. oryzae Mo28S* normalized to rice *Ubiquitin1* gene. Bar plots in B represent mean values ± SEM from four biological replicates (each comprising 6–8 plants). Statistical significance was determined by two-way ANOVA followed by Tukey's multiple comparison test, comparing values within and between Pi and Pi+Phi treatments. Bars labeled with the same letter are not significantly different from one another.

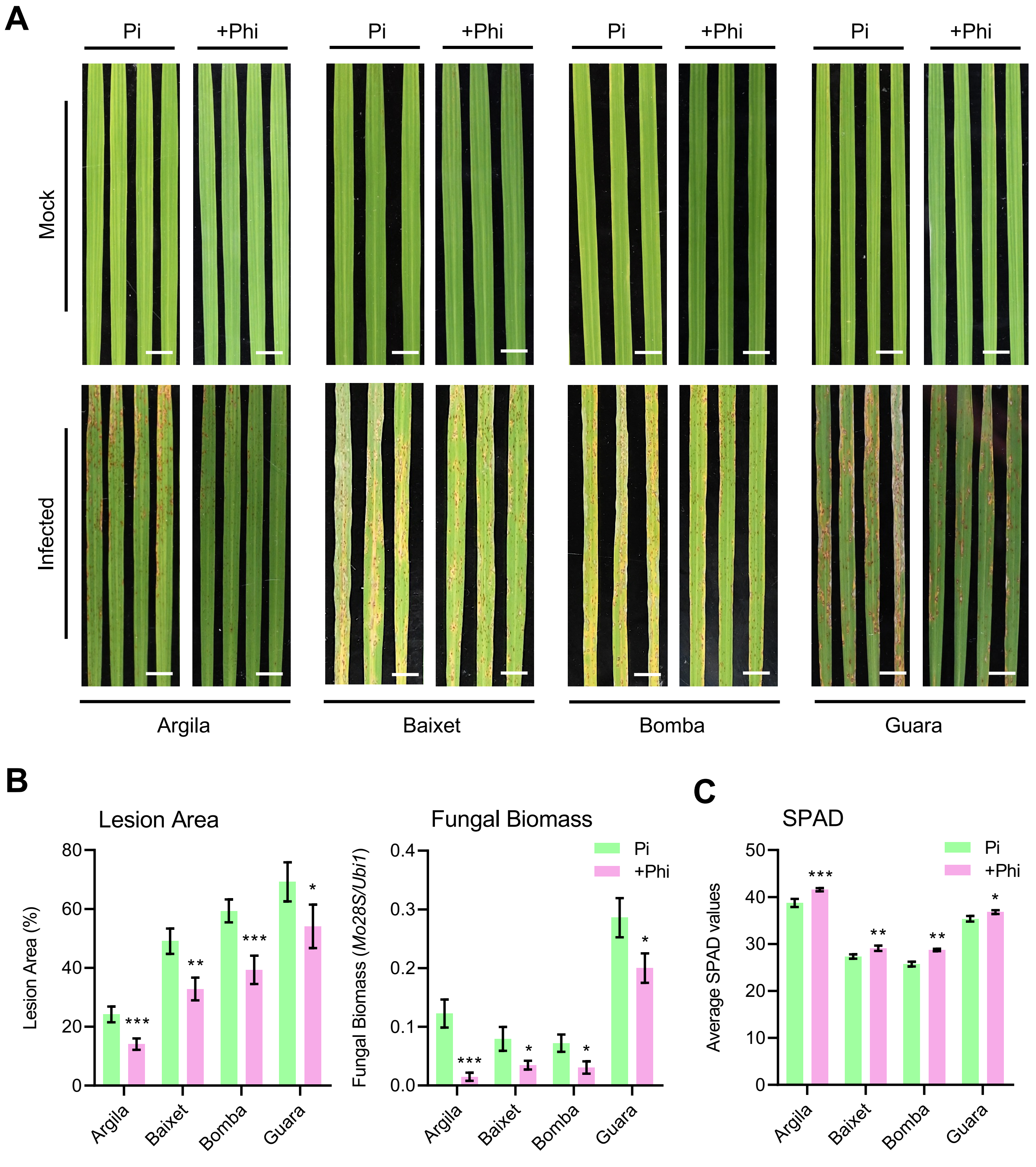


**Supplementary Figure S4.** Protective effect of Phi against *M. oryzae* infection in *japonica* rice cultivars grown under high Pi supply. Argila, Baixet, Bomba, and Guara cultivars were grown under 2.5 mM Pi for 2 weeks, without Phi supplementation or with Phi (0.25 mM Phi), and then mock-inoculated or inoculated with a suspension of *M. oryzae* spores (5 x $10^5$ spores/ml). Three independent experiments were carried out with similar results. (**A**) Representative images of blast symptoms in rice leaves. Images correspond to the 3rd leaf of plants, 7 days post infection. Scale bar = 1 cm. (**B**) Left panel, Lesion area of Phi-treated plants. Right panel, Fungal biomass quantified at 7 dpi by qPCR for *M. oryzae Mo28S* normalized to rice *Ubi1*. (**C**) SPAD measurements of the 3rd leaf. Bar plots represent mean values ± SEM from four biological replicates (each comprising 6 plants per pool). Statistical significance was tested using Student's *t*-test, comparing values between the control Pi and corresponding Phi treatments (*$p < 0.05$, **$p < 0.01$, ***$p < 0.001$).

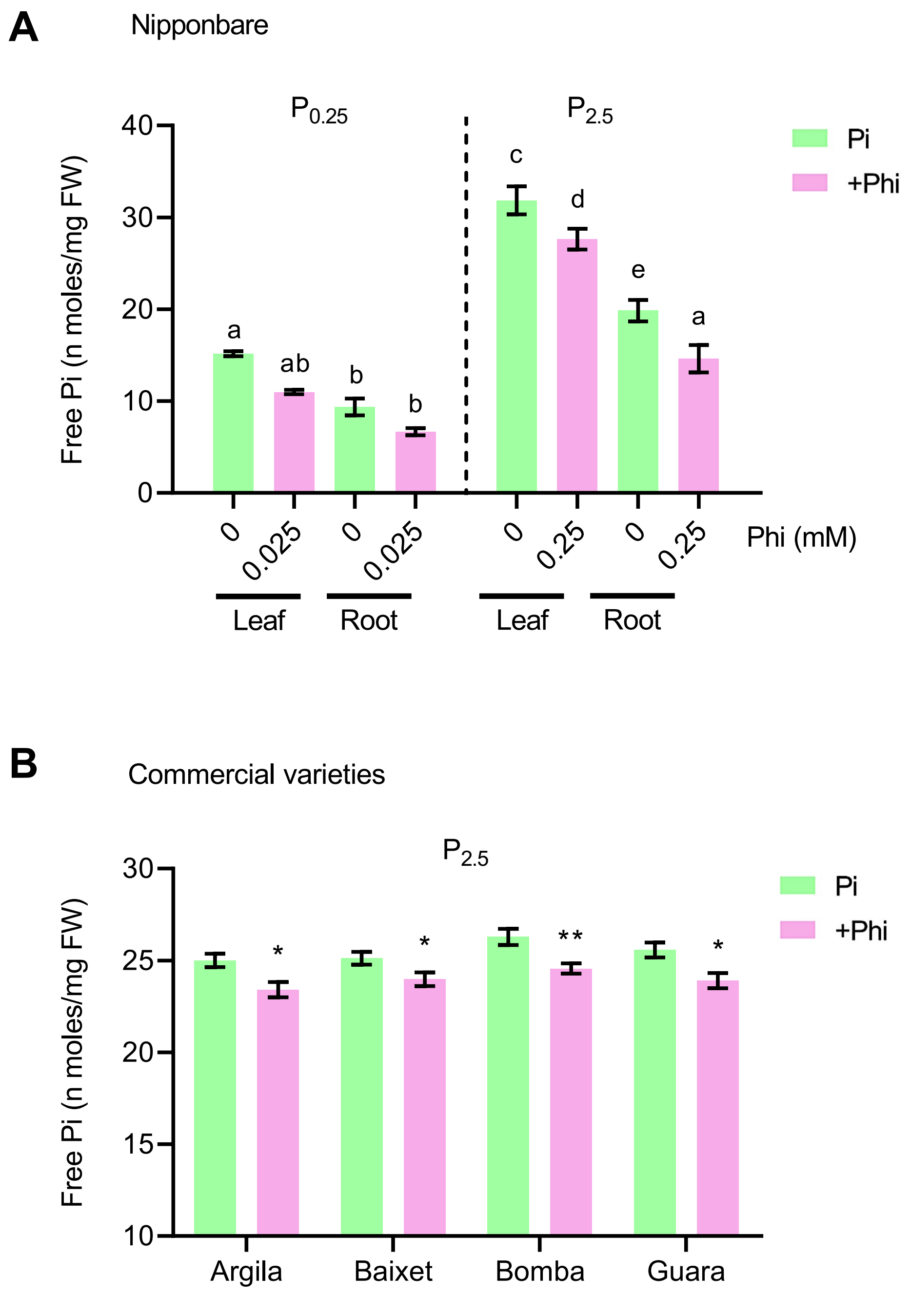


**Supplementary Figure S5.** Free Pi content in rice varieties supplemented, or not, with Phi. Rice plants were grown under optimal or high Pi supply. Leaf Pi content was determined from the 3rd leaf of Pi or Pi+Phi-treated plants. Root Pi content was assessed on entire roots. Data represents mean values and error bars ± SEM. (**A**) Free Pi content of Nipponbare plants; statistical significance tested by two-way ANOVA and Tukey's multiple comparison tests, within and between the Pi and Phi treatments and letters above each error bar represents significance, with same letters indicating no statistical difference. (**B**) Free Pi content of *Japonica* rice varieties; statistical significance was tested by Student's *t*-test compared to Pi only controls per condition, indicated by $*p < 0.05$, $**p < 0.01$.
